# Reconstructed Cell-Type Specific Rhythms in Human Brain link Alzheimer’s Pathology, Circadian Stress, and Ribosomal Disruption

**DOI:** 10.1101/2025.02.21.639499

**Authors:** Henry C. Hollis, Ashish Sharma, Patrick W. Sheehan, Leonard B. Maggi, Jason D. Weber, Jan A. Hammarlund, David A. Bennet, Vilas Menon, Erik S. Musiek, Ron C. Anafi

**Author notes:** Correspondence to: Dr. Erik Musiek, Dr. Ron Anafi. Contributed equally to this work (Henry C. Hollis, Ashish Sharma). Joint corresponding authors (Erik S. Musiek, Ron C. Anafi).

## Abstract

Alzheimer’s disease (AD) disrupts behavioral circadian rhythms, but its effects on molecular rhythms in the human brain are poorly understood. Using single-nucleus RNA sequencing from post-mortem cortical samples, we informatically estimated the relative circadian phases of 409 persons with and without AD dementia. We then reconstructed circadian expression profiles across cell types. While core clock rhythms were preserved in AD, many cell-type specific circadian outputs were disrupted. Rhythms in ribosomal biogenesis and oxidative phosphorylation were dampened across cell types. Similar losses in ribosomal gene expression rhythms were observed in APP/PS1 mice, which showed further reductions in ribosomal protein expression and polysome-mediated translation after circadian desynchrony. Exploratory computational modeling reveals that altered translation may contribute to the increased circadian variability seen in AD patients. These findings reveal altered cell-type specific circadian output rhythms in the brains of AD affected patients, and highlight disrupted ribosomal rhythms as a feature of AD.

## Introduction

Circadian, or daily, rhythms are everywhere in nature^1^, influencing many areas of behavior and physiology beyond cycles of sleep and wakefulness. Cell autonomous rhythms are driven by a core transcriptional-translational feedback loop involving activators CLOCK and BMAL1 along with inhibitors from the PERIOD and CRY family genes^2^. In mammals, the suprachiasmatic nucleus (SCN) coordinates these cellular rhythms across tissues^2,3^. Depending on the tissue, circadian time modulates the expression of hundreds or thousands of different genes and proteins, among which are disease genes and drug targets^4–6^.

AD is a neurodegenerative disease that accounts for nearly 60-80% of dementia cases^7^. Pathologically, the disease is characterized by amyloid-β (Aβ) plaques and tau associated neurofibrillary tangles (NFT). Symptomatically, AD is characterized by progressive memory loss and cognitive decline. However, circadian disturbance and sleep/wake disruption are also prominent features of the disease. Several studies of AD patients demonstrate fragmentation of behavioral circadian rhythms and phase delay^8,9^. Significant daily variability *within and between* AD patients is also a prominent feature of these data^10^. AD related circadian rhythm sleep disorders are particularly difficult to treat^9,11^. Moreover, these altered rhythms go beyond behavior, as AD patients demonstrate altered rhythms in core body temperature and hormone secretion^12–14^.

Circadian rhythmicity is regulated at the cellular level by conserved core clock genes, which control rhythmic gene expression in a cell- and tissue-specific manner. While studies have described alterations in core clock function and circadian gene expression patterns in AD patient-derived cells^15^, or in AD mouse models^16,17^, it is unknown how AD impacts circadian function at the cellular or molecular level in human brain.

Evidence suggests that the influence of AD on circadian rhythms is multilayered. Knockdown of genes critical to AD pathogenesis, including *APOE*, *PSEN1* and *PSEN2*, alter the expression of core clock genes and ultimately the period and amplitude of *in-vitro* rhythms^18^. At the level of neural circuits, AD targets neuronal populations involved in SCN synchronization^19,20^, along with downstream, orexinergic neurons regulating sleep and wake^21^.

Emerging data suggests that circadian dysfunction may, in turn, exacerbate AD pathology^22–24^. Circadian genes regulate redox balance and metabolism^25,26^, processes that are especially relevant to AD. In wild type mice, disruption of circadian function via deletion of the core clock protein BMAL1 in the hypothalamus or cortex results in astrogliosis, oxidative stress, and synaptic degeneration^27^ In an APP/PS1 mouse model of AD, local clock disruption via deletion of BMAL1 exacerbates plaque deposition. The clock appears to modulate both microglial mediated neuroinflammation^28,29^ and may influence Aβ synthesis^30^. Indeed, phosphorylated tau and Aβ both vary with the sleep wake cycle^22,31,32^.

Despite these basic science connections, translational circadian research in AD remains limited. Understanding how molecular rhythms change across cell types in the brains of AD patients could inform our understanding of AD pathology and reveal new therapeutic avenues.

## Results

### Assigning circadian phase using CYCLOPS allows circadian analysis of human AD postmortem transcriptome data

Transcriptional circadian rhythms are typically determined by euthanizing model organisms at regular intervals and measuring gene expression over time. As these organisms are well synchronized with each other and with the external environment, a sample’s collection time becomes an effective proxy for its internal circadian phase.

Ordering control (CTL) samples by collection time, molecular rhythms become apparent (Fig 1A, left). Obviously, this procedure is unethical in humans and as such researchers often rely on recorded time of death (TOD) from post-mortem expression data to reconstruct molecular rhythms^33,34^.

**Figure 1:**
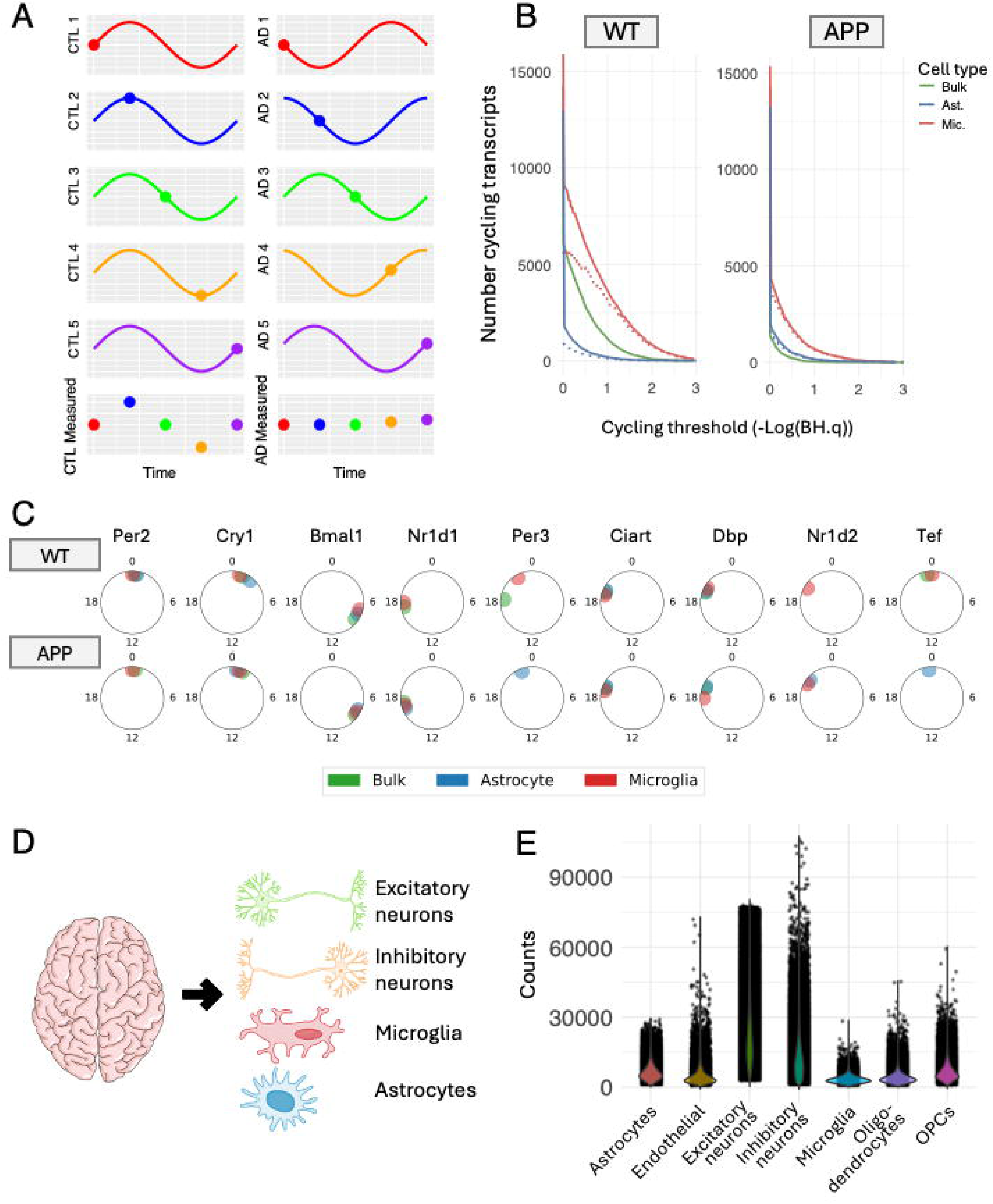
Inferring Circadian Phases of Aged CTL and AD dementia Post-Mortem Subjects. **(A)** *Schematic of the difficulty of identifying rhythms from AD population data.* Left: CTL subjects are well-entrained to the external environment and synchronized with each other. Rhythmic gene expression can be reconstructed plotting gene expression by TOD (left, bottom). Right: AD dementia subjects are poorly entrained to the external environment so that reconstructing rhythmic gene expression from TOD becomes unfeasible (right, bottom). **(B)** *Cell type-specific transcriptional rhythms are lost in bulk tissue analyses.* JTK analysis was performed on mouse cortical data with astrocyte specific (RiboTag) and microglia-specific (TRAP-seq) transcripts. *Left*: The number of cycling genes in WT bulk tissue (solid green), WT astrocytes (solid blue), and WT microglia (solid red) as a function of JTK cycling significance (-Log(BH.q)). Dashed lines indicate genes found to be cycling in astrocytes (dashed blue) or microglia (dashed red) but not in bulk tissue. *Right*: The same analysis repeated on APP mice data. **(C)** *Synchronization of the core clock across cortical cell types in WT and APP mice*. Radial plots display the acrophase (CT) of core clock genes across cell types. Top: WT mice; Bottom: APP mice. Clock genes in bulk tissue (green), astrocytes (blue), and microglia (red) appear synchronized in WT and APP. **(D)** *Schematic of single-nucleus sequencing aggregation by subject and cell type*. Aggregation reduces noise and facilitates analysis of transcriptional rhythms. **(E)** *Excitatory neurons contribute most of the RNA counts in the ROSMAP data.* Horizontal axis: Cell types in ROSMAP data. Vertical axis: Total raw RNA counts per cell.

There are several problems with applying this approach to AD. First, existing brain expression datasets, such as the Accelerating Medicines Partnership – AD (AMP-AD), often do not include TOD or other traditional circadian phase markers (e.g., melatonin levels). Moreover, as AD patients are seldom observed at night, times of death may be subject to large error. Even with accurate TOD information however, TOD-based analyses of AD samples may fail. AD patients have highly variable phases of circadian entrainment. The assumption that AD subjects are well synchronized with each other and well entrained to the external environment is suspect. The time on an external clock may not reflect a subject’s internal circadian phase, compromising TOD-estimated rhythms (Fig 1A, right).

Instead of relying on TOD, we infer each sample’s molecular circadian phase using the Cyclic Ordering by Periodic Structure (CYCLOPS 2.0) algorithm^35,36^: Plotting the expression of one cycling gene against the expression of another out-of-phase gene forms an ellipse (Fig S1A-D). Plotting the expression of three cycling genes against each other, a 2-dimensional ellipse again arises. The expression of hundreds or thousands of circadian transcripts lies on a single ellipse in gene expression space. The relative position of each sample on the ellipse reflects its relative position in the circadian cycle. CYCLOPS uses an autoencoder to identify circular structure in high-dimensional data that includes many rhythmic molecules. CYCLOPS 2.0^36^ allows the autoencoder to adjust these data for non-circadian confounders such as sequencing batch and disease state.

We previously validated CYCLOPS’s ability to order bulk transcriptomic data derived from human cortical brain samples collected at autopsy^35^. However, because the brain is composed of a heterogenous mix of cells, we hypothesized that cell-specific rhythms would be obscured in bulk RNAseq data, and that single-cell data may be necessary to identify altered rhythms in specific cell types relevant to AD. To begin to examine this, we reanalyzed an existing circadian transcriptomic dataset collected at 2-hour intervals from wild type (WT) and APP/PS1-21 (APP) transgenic mice, consisting of bulk RNAseq data from mouse cortex^37^, as well as microglia- and astrocyte-specific gene expression data obtained via RiboTag and TRAP-Seq, respectively. Our analysis of data from both WT and APP mice reveals that transcriptional rhythms specific to individual cell types are often lost in bulk data (Fig 1B). These data suggest that the circadian signal for CYCLOPS ordering may be enhanced in cell-type specific data. Moreover, within both WT and APP mice, the different cell types of the brain appeared to be synchronized within a sample, sharing a common circadian phase (Fig 1C) so that an ordering based on cell types with strong circadian signal could be applied to other cell types with weaker signals.

To create a cell-type specific circadian transcriptomic atlas in the human brain, we reanalyzed recently published single-nucleus RNA sequencing data from 159 persons with AD dementia and 250 aged CTL from (ROSMAP)^38^ (Tab 1). Subject condition labels were based on the clinical consensus diagnosis of Alzheimer’s dementia at time of death (COGDX). All samples were obtained from the dorsolateral prefrontal cortex (DLPFC). Clustering and cell type labels were obtained from the original publication^39^.

Because single-nucleus data is often noisy, we sought to minimize noise by analyzing these data at the pseudo-bulk level, aggregating single-nuclei from cells of the same type in each sample (Fig 1D). Excitatory neurons contribute the most RNA counts (Fig 1E) in these ROSMAP data, so we first evaluated the circadian signal in those cells.

To optimize our analysis, we identified specific subclusters of excitatory neurons with the strongest circadian signal. The clock correlation distance (CCD) is a measure of core clock organization in unordered data^40^. CCD compares the gene-gene correlation matrix of 12 core clock genes with an established reference. As non-circadian variation grows, the circadian signal is obscured and the CCD increases. Using the pseudo-bulk counts generated by combining excitatory neuron subclusters 3 and 5 results in a correlation matrix that most strongly resembles the reference (Fig 2A-B). Notably, the separate gene-gene correlation matrices for excitatory subclusters 3 & 5 do not individually resemble the established reference (Fig S2A-B). Moreover, the quantity of exc.3 neurons per subject is negatively correlated to the quantity of exc.5 neurons per subject (r = -0.28) (Fig S2C). Thus, we suspect that the original clustering may have mislabeled biologically synonymous cells from different circadian times as being distinct cell types.

**Figure 2:**
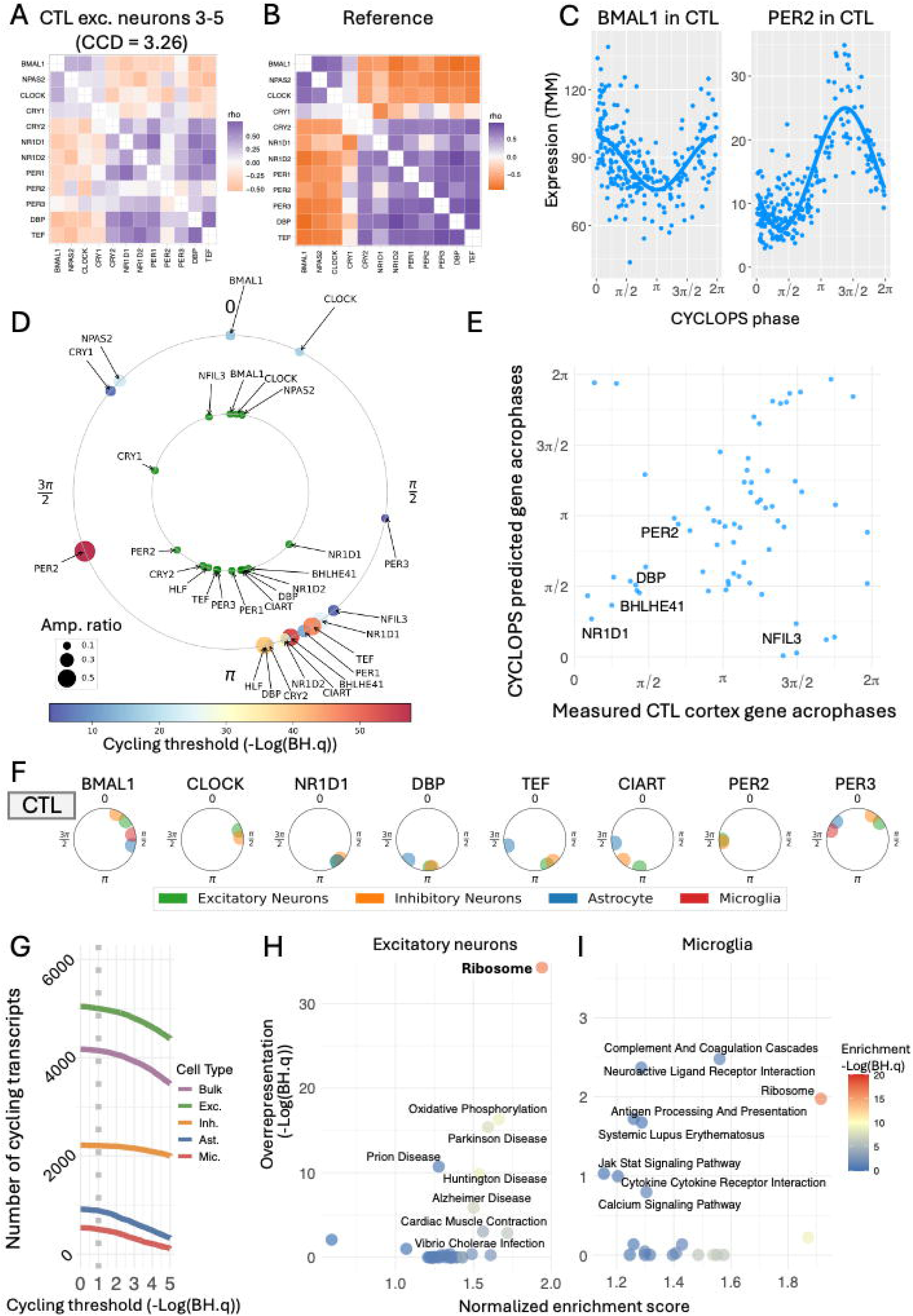
The CYCLOPS Ordering of CTL Samples Matches Biological Expectation. **(A)** *Correlation matrix of 12 core clock genes* in excitatory neuron subtypes 3-5 aggregated by subject closely resemble the reference matrix (CCD = 3.262) **(B)**, indicating a strong circadian signal in these neurons. These subtypes were used for informatic ordering. **(C)** *Gene expression as a function of CYCLOPS inferred circadian phase*. In CTL excitatory neuron subtypes 3-5, core clock genes BMAL1 and PER2 exhibit strong rhythmicity, consistent with expectation. **(D)** *Relative acrophases of core clock genes in CTL excitatory neuron subtypes 3-5 compared to biological expectation.* The inner ring (green dots) shows relative timing of core clock acrophases from the mouse atlas^4^. The outer ring shows the relative timing of core clock acrophases in CTL excitatory neuron subtypes 3-5. The color of dots on the outer ring corresponds to significance of cycling, and the size of the dots on the outer ring denotes amplitude ratio (amplitude / MESOR). **(E)** *Comparison of CYCLOPS predicted acrophases of clock outputs (full pseudo-bulk transcriptome) with prior bulk transcriptional data from human cortex.* Horizontal axis: Previously measured acrophases of cycling genes (BH.q < 0.2) in CTL human cortex^33^. Vertical axis: CYCLOPS-predicted acrophases of cycling genes (BH.q < 0.1 & amplitude/ MESOR ≥ 0.2) from CTL all cell pseudo-bulk data (aligned to measured acrophases). (Jammalamadka ranked circular correlation = 0.39). **(F)** S*ynchronization of the core clock across cell types in aged CTL human cortex.* Radial plots display the CYCLOPS-predicted acrophases of core clock genes in aged CTL humans. Colored dots represent acrophases in excitatory neurons (green), inhibitory neurons (orange), astrocytes (blue), and microglia (red). Only core clock transcripts with cycling significance BH.q < 0.1 in a given cell type are shown. **(G)** *Number of cycling genes across cell types.* Cycling genes were identified using cosinor regression. The number of cycling genes is shown for CTL all-cell-pseudo-bulk (purple), excitatory neurons (green), inhibitory neurons (orange), astrocytes (blue), and microglia (red) as a function of statistical cycling threshold (BH.q value). Vertical dashed line indicates cycling BH.q = 0.1. Gene quantities are shown for genes with amplitude ratio ≥ 0.2. **(H-I)** *Pathway-level cycling analysis in CTL excitatory neurons* **(H)** *and microglia* **(I)**. Two complementary approaches were used to identify genes sets and pathways strongly associated with cycling genes: enrichment analysis using fGSEA and overrepresentation analysis using Enrichr. Horizontal axis: fGSEA normalized enrichment score. Vertical axis: Significance of overrepresentation (-Log(BH.q)) from Enrichr. Color indicates fGSEA enrichment significance (-Log(BH.q)).

### The CYCLOPS ordering of CTL samples matches biological expectation

We applied CYCLOPS 2.0 to cell type-specific pseudo-bulk data from combined excitatory neuron subclusters 3-5, treating sequencing batch and disease status as covariates. The ordering passed established quality metrics (stat error p<0.05, smoothness <1)^35,36^. Notably, we also used CHIRAL^6^, an alternative algorithm, to order these same data. The two orderings were well correlated (Jammalamadaka ranked circular cor=0.84, Fig S2D). We initially focused on CTL subjects, performing cosinor regression to identify rhythmic transcripts in the neuronal 3-5 subcluster, requiring an FDR < 0.1 and a relative amplitude = amplitude/Midline-estimating statistic of rhythm (MESOR) ≥ 0.20. This regression model accounted for non-circadian variables including sex, post-mortem interval, and sequencing batch. Examples of cycling core clock genes in excitatory neuron subtypes 3-5 are shown in Figure 2C. The relative acrophases of core clock genes are well conserved across mammals and provide an important biological check on circadian reconstructions. The relative acrophases of core clock genes in the 3-5 subcluster estimated by CYCLOPS closely align with the acrophases of these genes in mice^4^ (Fig 2D).

We used the circadian phases determined above to identify cycling transcripts in the full pseudo-bulk transcriptome (including all cell types). The CYCLOPS-estimated acrophases from these pseudo-bulk DLPFC samples generally agreed with acrophases previously measured in bulk BA11 human cortex^33^ (Fig 2E). We next analyzed cell type-specific pseudo-bulk data (excitatory neurons, inhibitory neurons, astrocytes, and microglia). Using the sample phases estimated from excitatory neurons 3-5 we identified cell type-specific rhythms. Consistent with our mouse data, the rhythms of core clock genes were generally synchronized across these cell types (Fig 2F). Applying cosinor regression and using the same thresholds (Fig 2G), we identified 5001 cycling transcripts in excitatory neurons, 2209 in inhibitory neurons, 885 in astrocytes, and 504 in microglia.

### Many transcripts involved in pathways implicated in neurodegenerative disease show cell-type specific rhythms in CTL human brains

Two complementary approaches, enrichment (fGSEA)^41^ and overrepresentation (Enrichr)^42–44^, were used to identify pathways cycling in CTL subjects, separately analyzing data from excitatory neurons (Fig 2H), inhibitory neurons (Fig S3A), microglia (Fig 2I), and astrocytes (Fig S3B). Focusing on neurons and looking at the results of both methods, gene sets describing pathways for neurodegenerative diseases (including Parkinson, Huntington, Alzheimer, and Prion disease pathways) were among those that showed the strongest enrichment for cycling. Notably, many of the genes enriched in multiple of these pathways encode mitochondrial electron transport chain components (NDUFA13, NDUFA11, COX6A1, COX6B1, UQCRB), proteasome subunits (PSMB6, PSMD8), and cytoskeletal elements (TUBB3, TUBB6, TUBA1A, TUBA1B, TUBA4A), suggesting circadian regulation of cellular homeostasis processes relevant to neurodegeneration. These findings indicate that key neurodegeneration-associated pathways exhibit circadian rhythmicity in healthy individuals. In microglia, pathways related to complement activation and JAK-STAT signaling were enriched for circadian rhythmicity. This may be particularly relevant, as microglial complement expression is implicated in synapse loss in AD models^45,46^, while increased JAK-STAT signaling occurs early in AD pathogenesis^47^ and offers a potential therapeutic target^48^. Astrocytes demonstrated enrichment for cycling of type 2 diabetes (HKDC1, IRS2, GCK) and glucose pathways. Both microglia and astrocytes showed enrichment for cycling genes related to antigen presentation, suggesting a possible circadian basis to glial-mediated recruitment of T cells into the brain, a process which has been strongly implicated in AD-related neurodegeneration^49^. Notably, oxidative phosphorylation and ribosome biogenesis pathways demonstrated a high degree of circadian coordination across multiple cell types (Figs 2H, 2I, and Fig S3).

### The core molecular clock is relatively unaffected by AD

We next used our CYCLOPS derived ordering to analyze data from AD affected subjects. In addition to using cosinor regression to identify rhythmic transcripts, we used nested models to identify rhythms that significantly differed between CTL and AD dementia subjects^35,50,51^. These include differences in amplitude or acrophase.

Moreover, the nested models allowed us to distinguish between differences in a gene’s rhythmic profile and its rhythm adjusted mean expression (MESOR).

Using the CYCLOPS derived sample phases to identify rhythmic genes in excitatory neuron subclusters 3-5 in AD patients, we again find that the relative ordering of genes in the core clock matched biological expectation (Fig 3A). Focusing on the expression profiles of these genes in both the CTL and AD populations revealed no significant differences in the reconstructed circadian profiles (Fig 3B). Comparing core clock rhythms in CTL and AD dementia cases across all cell types, we found almost no differences. The single exception was a greater amplitude of TEF cycling in the microglia of AD dementia subjects. As in CTLs, core clock gene rhythms appeared to be synchronized across cell types in AD subjects (Fig 3C). The robustness of the core clock in human AD patients mirrors our previous findings comparing APP and WT mice^37^.

**Figure 3:**
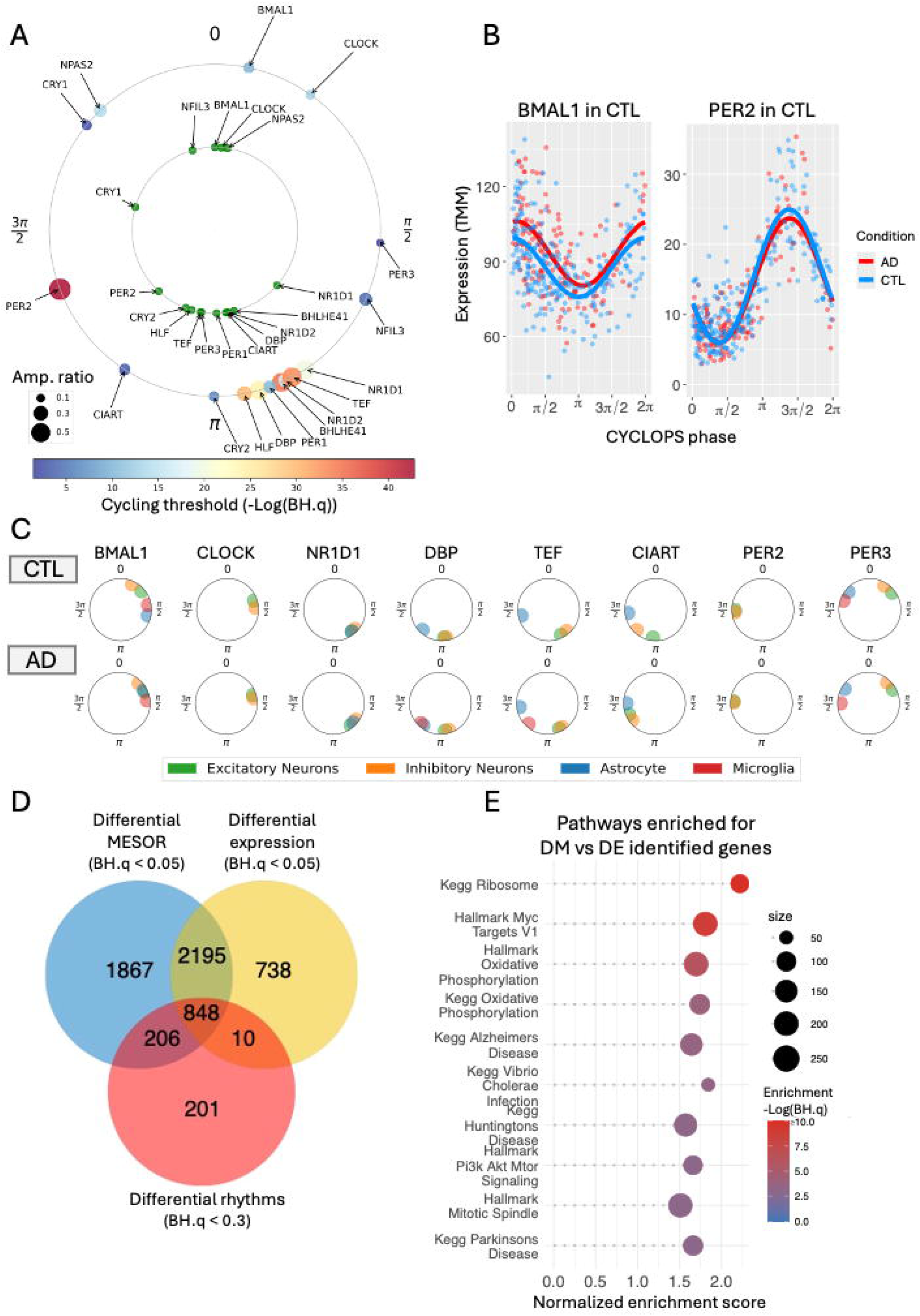
The Core Molecular Clock is Robust to AD. **(A)** *Relative acrophases of core clock genes in AD excitatory neuron subtypes 3-5 compared to biological expectation.* The inner ring (green dots) shows relative timing of core clock acrophases from the mouse atlas. The outer ring shows the relative timing of core clock acrophases in AD excitatory neuron subtypes 3-5. The color of dots on the outer ring corresponds to significance of cycling, and the size of the dots on the outer ring denotes amplitude ratio (amplitude / MESOR). **(B)** *Gene expression rhythms of BMAL1 and PER2 remain strong in AD.* CYCLOPS-inferred circadian phases reveal robust rhythms for BMAL1 and PER2 in both CTL (blue) and AD (red) excitatory neuron subtypes 3-5, with no significant rhythm differences. **(C)** *Across cell types, the synchronization of core clock transcripts is preserved in AD.* Radial plots display CYCLOPS-predicted acrophases of core clock genes across cell types in aged CTL (top, repeated from Figure 2F for comparison) and AD (bottom) affected humans. Core clock rhythms remain synchronized across excitatory neurons (green), inhibitory neurons (orange), astrocytes (blue), and microglia (red) in AD. **(D)** *Circadian informed analysis provides unique insights.* The Venn diagram illustrates overlapping gene sets in excitatory neurons: genes with significantly different MESOR in CTL and AD (blue circle, BH.q < 0.05), differentially expressed genes in CTL and AD using standard analysis (yellow circle, BH.q < 0.05, edgeR), and genes with significantly differential rhythms in CTL and AD (red circle, BH.q < 0.3, testing cycling genes only with BH.q < 0.1 & amplitude/MESOR ≥ 0.2). **(E)** *Analysis of pathway level differences in AD subjects likely to be missed using standard DE.* Genes were ranked comparing our ability to detect AD related changes using either DM or standard DE methods (log p value ratio). Gene sets enriched for genes more easily identified by DM are shown. Vertical axis: KEGG and Hallmark pathways. Horizontal axis: fGSEA normalized enrichment score. Dot size represents number of genes within a pathway, and color represents enrichment significance.

Importantly this circadian analysis provides unique AD related insights. We performed differential expression (DE) analysis with edgeR^42–44^ to identify cell-type specific psuedobulk transcriptional changes between CTL and AD subjects. We compared these DE genes with those identified through differential rhythmicity (DR) and differential MESOR (DM) tests. A Venn diagram describing the overlap of these results in excitatory neurons is shown in Figure 3D. While many genes are identified by 2 or 3 methods, others are only identified through circadian informed analysis. When we ranked genes by our relative ability to identify transcriptional changes in AD subjects using standard DE and circadian adjusted DM methods, we find that genes belonging to rhythmic pathways would be more easily missed by standard DE methods (Fig 3E).

### Circadian adjusted data clarifies expression changes in known AD risk genes

In order to further understand how circadian rhythms might impact AD-related gene expression, we next focused on a compilation of genes implicated in GWAS studies for variants that incur added risk for late onset AD^52^. Many of these transcripts met our criteria for circadian rhythmicity (BH.q <0.1, amplitude / MESOR ≥0.2) in both AD and CTL subjects (Fig 4A). Notably, the canonical AD risk gene APOE exhibited rhythmic expression in CTL astrocytes and microglia, while other key AD genes (APP, PSEN1, PSEN2) showed weaker rhythmicity across CTL cell types (BH.q < 0.1 but amplitude / MESOR < 0.2). Other AD risk genes, including COX7C a gene important in mitochondrial respiration and SEC61G involved in the transport of proteins across the endoplasmic reticulum, showed significantly dampened rhythms comparing AD dementia cases relative to CTL (Fig 4B-C).

**Figure 4:**
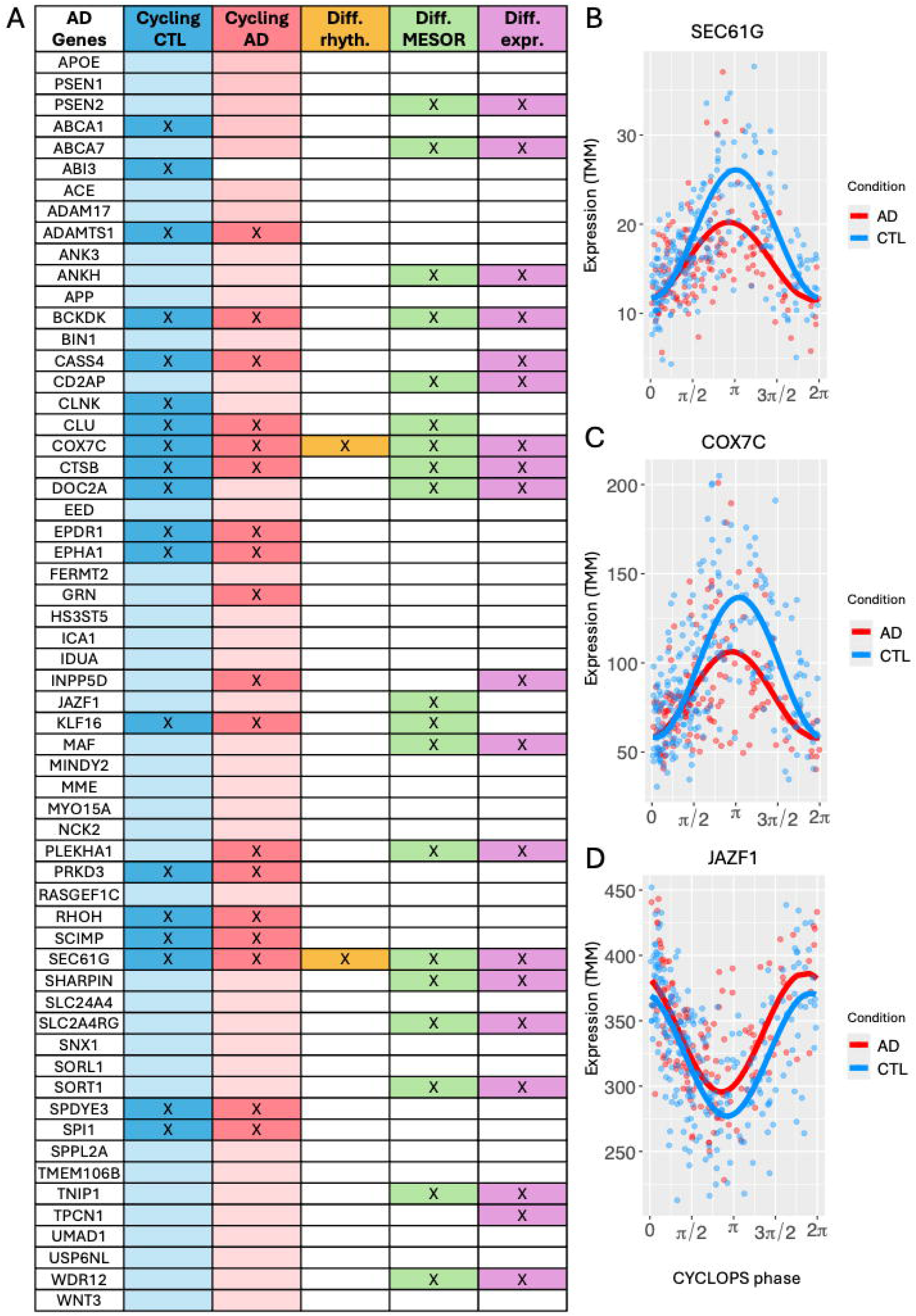
Circadian adjusted data clarifies expression changes in known Alzheimer’s Disease risk genes. **(A)** *Circadian analysis of known AD risk genes.* An “X” in the *Cycling in CTL/AD* columns indicates genes that meet the criteria for rhythmicity (BH.q <0.1 & amplitude/MESOR ≥ 0.2). Lightly shaded boxes in these columns, without an “X”, indicate weaker rhythms (BH.q <0.1 but amplitude/MESOR < 0.2). An “X” in the *Differential rhythms* column marks genes with significantly altered rhythms between CTL and AD (BH.q < 0.3). An “X” in the *Differential MESOR* column denotes significant basal expression differences between CTL and AD, controlling for circadian variation (BH.q<0.1). An “X” in the *Differential expression* column indicates significant differential expression (edgeR, BH.q < 0.1). **(B-D)** *Rhythmic reconstruction of key AD risk genes.* Expression of SEC61G **(B)**, COX7C **(C), and** JAZF1 **(D)** are shown as a function of CYCLOPS-inferred phase in CTL (blue) and AD (red).

As might be expected, several genes with known AD risk variants also showed evidence of differential expression comparing subjects with and without AD. Interestingly, differential MESOR analysis identified genes (CLU, KLF16, JAZF1) with basal expression differences that standard differential expression analysis overlooked (Fig 4A). The CLU gene encodes clusterin, a secreted protein involved in modulating both amyloid-beta and tau pathology^53,54^, while JAZF1 (Fig 4D) plays a crucial role in regulating ribosome biogenesis and aminoacyl-tRNA synthetase homeostasis^55^.

### Ribosome and oxidative phosphorylation pathways exhibit differential cycling in AD

We used both fGSEA and Enrichr to identify pathways associated with differentially rhythmic genes. Across cell types, differentially rhythmic genes were consistently enriched and overrepresented for the KEGG ribosome and oxidative phosphorylation pathways (Fig 5A-C). In contrast, other pathways, such as spliceosome regulation, exhibited cell-type-specific changes. Among the ribosomal transcripts demonstrating altered rhythmicity in AD, almost all showed reduced amplitude. Heat maps depicting rhythmic ribosomal transcripts in both CTL and AD subjects in microglia are shown in Fig 5D. Heat maps for neurons and astrocytes show a similar pattern (Fig S4).

**Figure 5:**
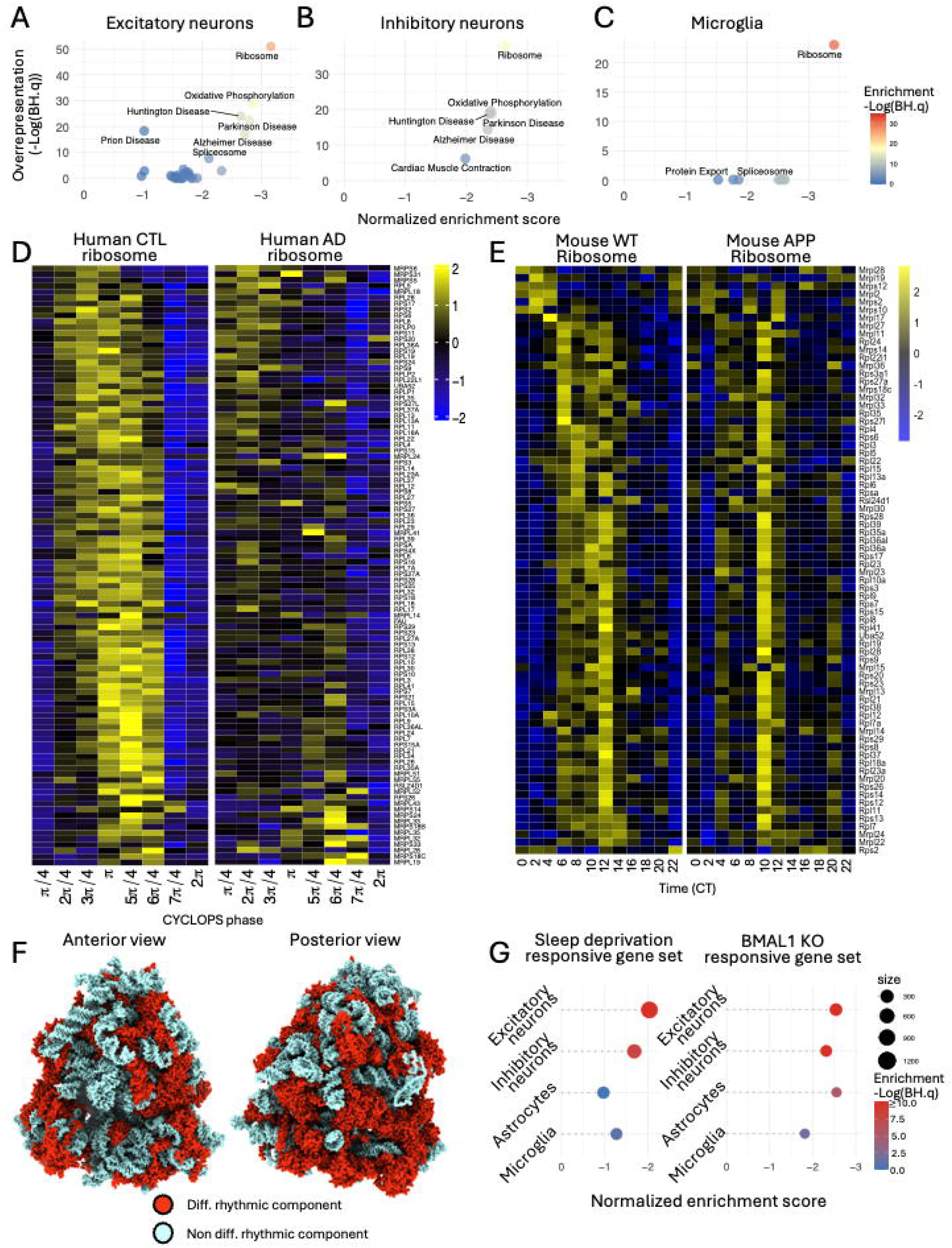
Ribosome and Oxidative Phosphorylation Pathways are Associated with Differentially Cycling Genes in AD. **(A-C)** *Pathways associated with differentially rhythmic genes* in excitatory **(A)**, inhibitory neurons **(B)**, and microglia **(C)**. Two complementary approaches were applied: enrichment analysis using fGSEA and overrepresentation analysis using Enrichr. Horizontal axis: fGSEA normalized enrichment score. Vertical axis: Significance of overrepresentation (-Log(BH.q)) from Enrichr. Color indicates fGSEA enrichment significance (-Log(BH.q)). For fGSEA analayis, genes were ranked by Log(amplitude AD/ amplitude CTL) so negative enrichment scores correspond to pathways associated with genes that lose amplitude in AD. The thresholds for significant rhythmicity in neuronal cell types were BH.q < 0.1 & amplitude/MESOR ≥ 0.2. The amplitude ratio threshold was relaxed to amplitude/MESOR ≥ 0.1 for microglia, which had fewer nuclei and sparser data. **(D)** *Circadian heatmap of ribosome genes in human microglia.* Heatmaps show cycling ribosome genes in CTL (left) and AD (right) microglia. Rows are ordered by CYCLOPS-predicted acrophases, and columns represent binned CYCLOPS phases. **(E)** *Circadian heatmap of ribosome genes in WT and APP/PS1 microglia.* Heatmaps display cycling ribosome genes from WT (left) and APP/PS1 (right) microglia, with rows sorted by acrophases and columns showing binned time (CT). **(F)** *Differentially rhythmic ribosomal proteins in excitatory neurons,* illustrated on structure of the human 80S ribosome^82^. Protein components colored with ChimeraX^83^. **(G)** *Enrichment analysis of sleep deprivation and BMAL1 KO responsive pathways.* Genes that were differentially rhythmic in the brains of AD patients were tested for enrichment with regards to genes that are responsive to sleep deprivation in murine cortex (left, differential expression FDR < 0.01) or genes responsive to BMAL1 knockout in murine midbrain (right, differential expression BH.q < 0.1). Genes were ranked by Log(amplitude AD / amplitude CTL) in each cell type, and fGSEA was used to test for enrichment using the specified gene sets. A negative enrichment score indicates pathways associated with genes losing rhythmic amplitude in AD. The vertical axis represents human cell types, the horizontal axis shows normalized enrichment scores, dot color reflects fGSEA significance, and dot size indicates the number of genes in the pathway.

Supporting these human data, we again examined the published mouse microglial Ribo-Tag-seq dataset^37^ and observed rhythmicity of KEGG Ribosome pathway transcripts in microglia from WT mice, while these rhythms were blunted in plaque-bearing APP/PS1 mice (Fig 5E). As illustrated in Figure 5F, the reduced rhythmicity observed in AD patients affects a large fraction of the ribosomal structure in excitatory neurons.

Ribosomal transcripts did not simply show a reduction in amplitude in AD subjects, but also a reduction in total expression. Using differential MESOR analysis to account for circadian time, we found that AD is broadly associated with a reduction in the rhythm corrected mean expression in ribosomal and oxidative phosphorylation related transcripts (Fig S5).

While our primary analysis stratified samples by clinical diagnosis (COGDX), we also examined differences based on neuropathological CERAD and Braak scores. CERAD scores reflect neocortical neuritic plaque density, whereas Braak scores measure paired helical filament (PHF)-tau tangles. The reduced rhythmicity of ribosomal genes in AD dementia cases was more strongly associated with β-amyloid plaque load (CERAD) than with PHF-tau tangles (Braak). We found that genes that were differentially rhythmic as a function of CERAD score were enriched for the ribosome pathway in microglia, inhibitory neurons, and astrocytes. Identifying transcripts that were differentially rhythmic as a function of Braak score did not show an association with the ribosome pathway (Fig S6A-D).

Metabolomics data from a subset of these cortical samples has been previously published^56^. Using the circadian phases assigned to each sample based on gene expression, we assessed cycling and differentially cycling of metabolites (Fig S6E-G). Given the relatively limited number of our samples profiled with metabolomics, we used the MetaboAnalyst platform to integrate our metabolomic and transcriptional data and identify altered rhythms in metabolic pathways. Circadian changes in metabolomic pathways were concentrated in energy generation (e.g., glycolysis and TCA cycle).

### Differentially cycling transcripts in AD are enriched for sleep and circadian stress response pathways

Previous mouse experiments have identified transcriptomic signatures responsive to sleep and sleep deprivation in the murine cortex^57^. Similarly, prior work has identified genes differentially expressed in the setting of disruption of the molecular circadian clock caused by *Bmal1* deletion in mouse midbrain tissue^58^. These studies have emphasized changes in protein translation, a key ribosomal function, and the unfolded protein response resulting from these interventions.

Sleep and circadian disruption are symptoms of AD dementia. We ranked transcripts in our human study by the loss of circadian amplitude comparing AD dementia cases and CTL. We then used custom gene sets describing the responses to sleep deprivation and circadian disruption for fGSEA analysis. In neurons, genes losing amplitude in AD were enriched for sleep deprivation response genes (Fig. 5G, left). Across all cell types, transcripts losing rhythmicity in AD were enriched for genes responding to molecular circadian disruption (Fig. 5G, right). Notably, pathway analysis of genes responsive to midbrain *Bmal1* deletion showed KEGG ribosome and oxidative phosphorylation pathways as the top hits^58^. Collectively, these results suggest that rhythmic transcriptional differences in AD subjects may involve a sleep or circadian stress component.

### Overlapping effects of APP/PS1 genotype and circadian desynchrony on ribosomal disruption in mice

Acknowledging that both AD pathology and AD related circadian disruption might contribute to the altered expression of ribosomal transcripts in AD patients, we aimed to experimentally disentangle these influences using a mouse model. Unlike human AD patients, the APP^swe^/PS1^ΔE9^ mouse model of amyloid plaque pathology (referred to as PSAPP) exhibits only modest circadian disruption early in life^16^. Thus, we studied PSAPP mice and non-carrier/wild-type (WT) littermate controls in three experimental cohorts: WT mice maintained in a 12:12 light-dark environment (WT+LD), PSAPP transgenic mice housed in a 12:12 light-dark environment (APP+LD), and PSAPP mice subjected to chronic jet lag protocol (APP+CJL), which consisted of repeated phase advances caused by shifting the lights-on 6 hours earlier each week. This protocol mimics jet lag caused by eastward travel, causing repeated desynchrony between the internal circadian system and the external lighting, without disrupting sleep or increasing cortisol levels^59,60^. We used 12-month-old WT and PSAPP mice in this study, as mice have stable, moderate plaque pathology at this age, and subjected them to LD or CJL for 12 weeks, then harvested mice after a one-week re-entrainment period, at a consistent time of day (ZT 4-6). This paradigm combines circadian stress with amyloid plaque pathology to model the circadian disruption commonly observed in human AD patients (Fig 6A).

**Figure 6:**
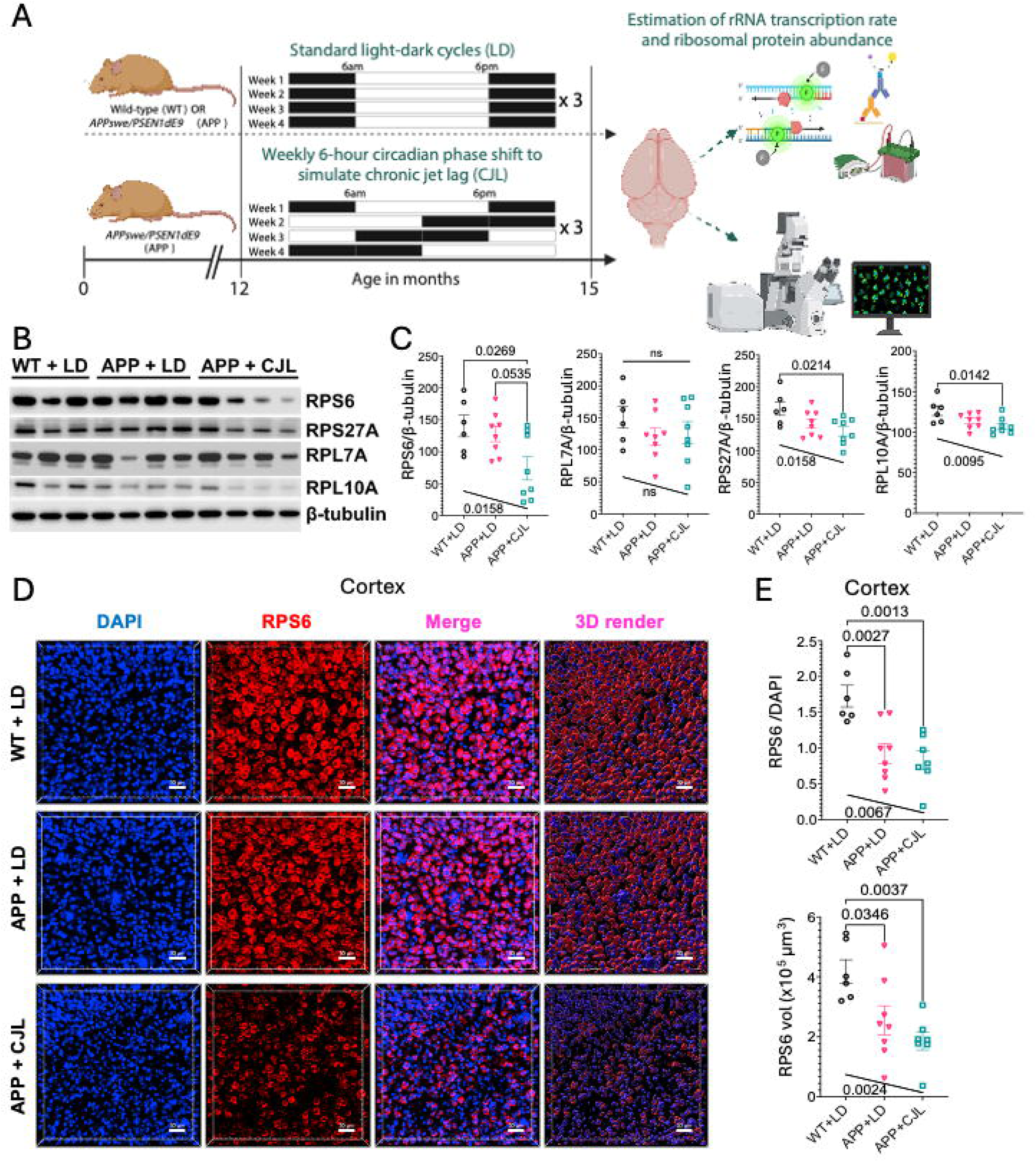
Overlapping effects of APP genotype and circadian stress on ribosomal disruption in mice. **(A)** At age 12 months, groups of APP^swe^/PS1^ΔE9^ AD model mice were and WT littermate controls were transferred to 12:12 controlled Light: Dark environment. At the same age, a third group of APP^swe^/PS1^ΔE9^ AD were subjected to chronic jet lag (CJL) protocol where the lighting schedule was advanced by 6 hours each week. N=6-8 mice were included in each group. After 3 months, all mice were given 1 week to fully entrain to a 12:12 light schedule before tissue was collected between ZT5 and ZT7. Cortical samples were sent for protein quantification by western blot and immunohistochemistry. Polysome profiling was employed to assess active translation. **(B)** *Western blots* of RPS6, RPS27A, RPL7A, RPL10A in WT+LD, APP+LD, and APP+CJL groups. **(C)** *Quantification of Western blots, protein abundance normalized by β-tubulin control.* Error bars indicate standard error of the mean (SEM). Significance values at the top (on horizontal lines) denote Tukey’s multiple comparisons test. Significance values near the bottom, by slanted lines, indicate significance of Kendall’s ranked correlation coefficient for ordinal association. **(D)** *Immunostaining of cortical slices* for RPS6 across experimental groups. **(E)** *Quantification of RPS6 immunostaining in cortex.* Error bars indicate standard error of the mean (SEM). Significance values at the top (on horizontal lines) denote Tukey’s multiple comparisons test. Significance values near the bottom, by slanted lines, indicate significance of Kendall’s ranked correlation coefficient.

Because the ribosome pathway was so consistently enriched for cycling genes across human cell types and in mouse brain, and lost rhythmicity in human AD and APP/PS1 mice, we focused our analysis on ribosomes. Ribosomes are nucleoproteins composed of various ribosomal-RNA (rRNA) species and ribosomal proteins. We analyzed elements of each structural component separately in all three experimental groups.

First, we assessed the transcription rate of *m47S* rRNA using RT-PCR and observed no significant difference, suggesting no change in the generation of structural rRNAs that give rise to ribosomes (Fig S7A). Next, we selected four ribosomal proteins which were found to be dysregulated in the human AD data. Two of the selected ribosomal proteins (RPS6 and RPS27A) are structural components of the small subunit (40S), and the other two (RPL7A and RPL10A) are components of the large subunit (60S). The abundance of these proteins in mouse cortex was quantified by Western Blot (Fig 6 B-C). Three of the four ribosomal proteins we examined (RPS6, RPS27A, and RPL10A) exhibited a statistically significant (p<0.05) reduction comparing the WT+LD and APP+CJL groups, while amyloid pathology alone did not cause statistically significant reduction. These three ribosomal proteins demonstrated a common trend with a stepwise reduction in abundance comparing the APP to WT cohorts and again with the addition of CJL to the APP genotype. Testing for an ordinal association between study group and protein abundance, using both parametric and non-parametric tests, this trend was statistically significant (p<0.05) for RPS6, RPS27A, and RPL10A. We further confirmed suppression of RPS6 protein expression specifically in the APP+CJL group via immunofluorescence microscopy using hippocampal and cortical sections (Fig 6D-E and Fig S7B-C). Significant reductions in RPS6 expression, which was primarily peri-nuclear, was observed in PSAPP+CJL compared to WT+LD in both regions. We again confirmed an ordinal association between study group and protein abundance, using both parametric and non-parametric tests, suggesting an additive effect of CJL on the APP genotype. Collectively, the data suggest that the additive effects of CJL and the APP genotype on ribosomes are mediated by changes in ribosomal protein abundance rather than alterations in rRNA transcription. Combining circadian stress with amyloid pathology, as can occur in human AD, leads to suppression of ribosomal protein expression in the mouse brain, similar to that observed at the transcriptional level in human AD.

### Amyloid pathology and circadian disruption combine to alter ribosome function

To investigate the impact of ribosomal protein suppression on protein translation, a fundamental function of ribosomes, we performed polysome profiling on mouse brain cortices (Fig 7A). This approach involves the separation and quantification of mRNA species based on ribosome occupancy using a sucrose gradient. The analysis revealed distinct peaks corresponding to mRNA species bound the 40S, 60S, and 80S ribosomal subunits, along with polysomes. The presence of polysome peaks indicates actively translating mRNAs bound by multiple ribosomes^61^.

**Figure 7:**
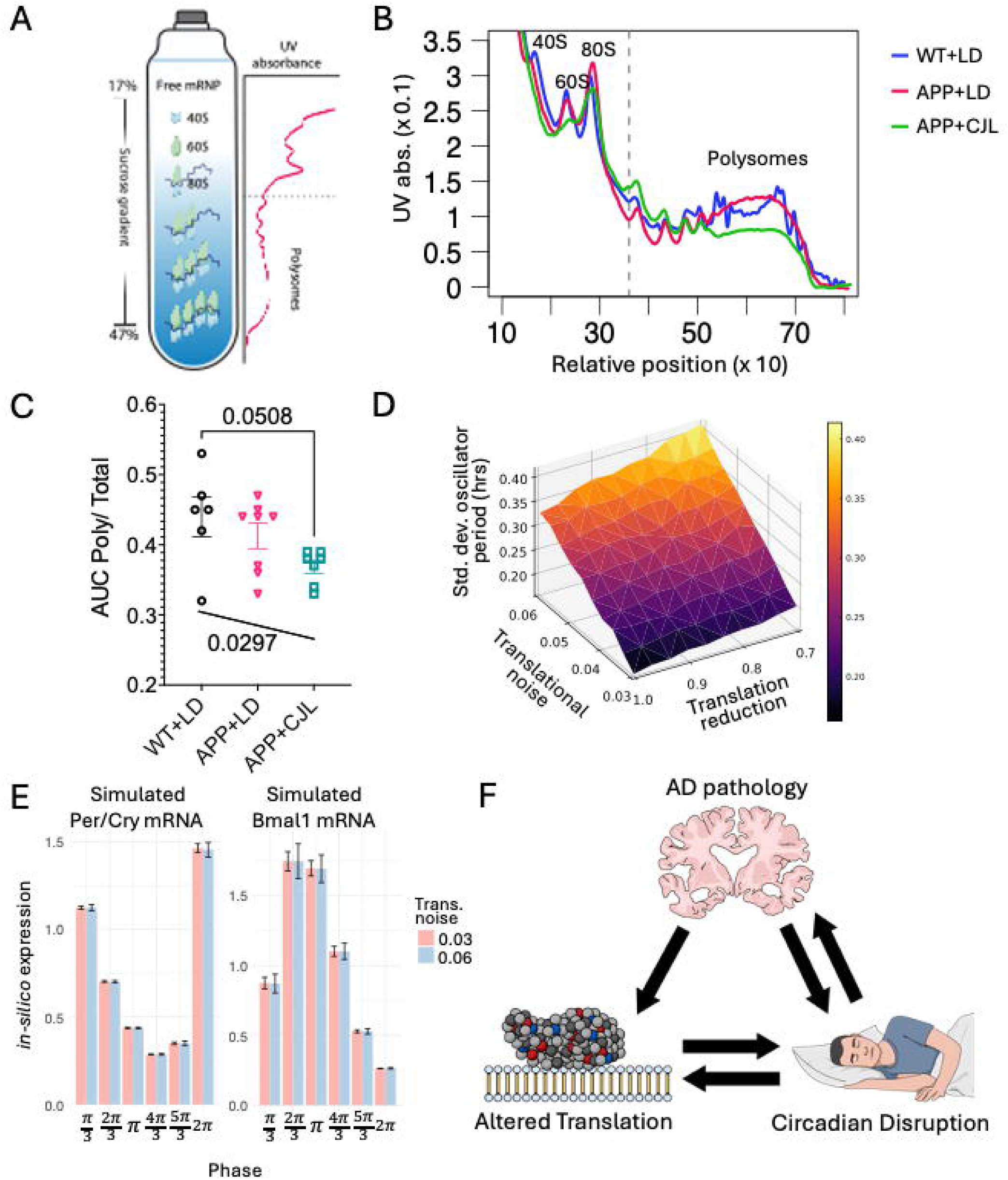
Functional consequences of reduced ribosomal components. **(A)** Schematic of polysome profiling. **(B)** Representative polysome profiling traces from WT+LD (blue) APP+LD (red) and APP+CJL (green) mice. **(C)** *Quantification of polysome profiling.* Area under the polysome peak is normalized by total area under the full absorption curve. **(D)** We added term describing translational stochasticity and a variable translation rate to a mathematical model of the circadian oscillator. Cycle-to-cycle circadian variability (SD of oscillator period) is plotted as a function of translational noise and translation rate. **(E)** Simulated Bmal1 and Per/Cry mRNA are not significantly changed by adding translational noise. **(F)** Schematic illustrating the tri-directional relationship between AD pathology, altered translation, and circadian disruption.

To assess translational activity, we aligned and analyzed polysome profiles using the QuAPPro tool^62^. Quantifying the area under the curve corresponding to polysomes and normalizing it to the total ribosome pool, we observed a significant reduction in polysome-associated mRNA in the APP+CJL group compared to the WT+LD group (Fig 7B-C). Both parametric and non-parametric tests for ordinal association demonstrate a significant negative association between experimental group and the normalized polysome signal. This finding suggests that circadian disruption adds to amyloid pathology in mice, further diminishing translational activity.

### Altered translation may affect the core clock oscillator

Given our findings of downregulated ribosomal components and altered protein translation in AD, we sought to investigate how such changes might influence the dynamics of the core circadian transcriptional-translational feedback loop. To explore this, we utilized a simplified, previously published mathematical model of the mammalian core clock oscillator^63^ which incorporates the transcription and translation of both positive and negative clock elements.

The initiation of protein translation is a stochastic process requiring the binding of a ribosome and related machinery to a specific RNA transcript. Previous studies suggest that decreased ribosome levels can lead to increased translational stochasticity^64,65^.

Other data suggests that reduced ribosomal abundance may slow the overall rate of translation^66,67^. We introduced Gaussian noise to the protein translation terms in the model along with a parameter describing a reduction in translation rate. We then tested the influence of both increased translational noise and reduced protein translation rates on oscillator function. The stochastic model showed cycle-to-cycle variation in period length. We systematically varied both parameters, conducting 20 independent simulations for various levels of translation reduction and translational noise. Our analysis revealed that the cycle-to-cycle variation in oscillator period length within each simulated subject (as assessed by the standard deviation of period lengths) increased as translation rate decreased (Fig 7D) and as noise increased. This finding indicates that the differences in ribosomal abundance observed in AD dementia cases have the potential to increase the variability of the core circadian clock period. Notably (Fig 7E) the altered dynamics in translation had minimal influence on the predicted core clock *transcript* expression consistent with our findings from human patients. This result is particularly intriguing in the context of AD, where patients are known to exhibit increased circadian variability in behavior and autonomic function^8,9,12–14^, aligning with our computational model’s predictions.

## Discussion

This study introduces a novel bioinformatic approach for identifying cell-type-specific molecular rhythms in aged human brains with and without Alzheimer’s disease. By surpassing the limitations of traditional TOD-based ordering^33,68^ and leveraging single-nucleus data^39^, our method provides unprecedented resolution in detecting rhythmic gene expression using previously generated transcriptomic data. In contrast to previous analyses of brain samples from subjects with and without AD related dementia, our approach reveals a more detailed and biologically relevant landscape of cell-type-specific rhythms. In control subjects, we found cell-type-specific cycling in numerous genes and biological pathways, including those linked to neurodegenerative diseases. In microglia, we uncovered strong enrichment of cycling transcripts in key AD-related pathways, such as complement activation, antigen processing, and JAK-STAT signaling—offering new insights that were previously obscured by methodologies that do not control for molecular clock time. In astrocytes and in bulk metabolomic analysis, we found that pathways involving glycolysis, energy utilization, and oxidative buffering were also strongly enriched for molecular rhythmicity.

Notably, despite well-documented behavioral circadian disruption in AD patients^12–14^, transcriptional rhythms in the core molecular clock remained largely unchanged. As in our previous study of WT and APP/PS1 mice^37^, core clock rhythms were generally synchronized across cell types in both AD patients and controls. However, AD patients exhibited significant disruptions in circadian outputs, including cell-type-specific changes in transcripts associated with AD risk. Most strikingly, we observed a broad reduction in the basal expression and cycling amplitude of transcripts related to ribosomal biogenesis and oxidative phosphorylation across cell types. Experimental validation confirmed these alterations in the microglia of AD model mice. Circadian analysis was critical in highlighting these changes: Circadian time accounts for substantial variance in these data, and many of these transcriptional changes are lost or muted using standard, untimed differential expression analysis.

Sleep disturbances and circadian stress, common in AD, have been linked to altered protein translation and increase oxidative phosphorylation^57,69–71^. We used the APP/PS1 model to disentangle the contributions of AD pathology and sleep/circadian stress on ribosomal protein loss. Three of four ribosomal proteins assayed (RPS6, RPS27A, and RPL10A) showed a stepwise decrease in abundance when circadian stress was added to amyloid pathology. Immunostaining confirmed significant reductions in RPS6 in both the cortex and dentate gyrus. These results demonstrate that some changes in the translational machinery associated with AD are intensified by circadian disruption.

Polysome profiling suggests that these reductions in ribosome abundance reduce the capacity for active protein translation.

Our results complement studies from the early 2000’s suggesting that ribosomal function is compromised in AD^72,73^. Using an *in-vitro* translation assay, Keller and colleagues found that ribosomes obtained from the brains of AD patients had decreased translational capacity and increased oxidative damage as compared to controls.

Focusing on the liver, Jouffe et al highlighted the importance of circadian control in the biogenesis of ribosomal components^74^. Many of the same ribosomal protein components found to cycle in the mouse liver also cycled in our human brain data.

Moreover, recent research in neurons suggests that individual ribosomal components are dynamically exchanged^75^. These changes in ribosomal composition may be a response to oxidative damage^75^ or may promote the preferential translation of specific classes of transcripts^76^. Given the impact of circadian rhythms on ribosomal dynamics, previous high-throughput studies that ignored circadian time may have underestimated these AD-related changes.

The additive effects of plaque pathology and circadian stress on ribosomal dysregulation are particularly relevant given the high prevalence of sleep and circadian disturbances in AD. This may explain why AD patients with pronounced circadian rhythm disruptions experience more rapid cognitive decline^77,78^. Theoretically, targeting sleep and circadian disturbances might mitigate ribosomal dysregulation and slow AD progression. This aligns with emerging evidence suggesting that circadian interventions, such as time-restricted feeding and bright light therapy, can improve AD pathology and symptoms^79,80^.

Altered protein translation has broad implications. Using computational modeling, we explored how reduced ribosomal abundance and altered protein translation might affect the core circadian oscillator. Translation initiation is stochastic and decreasing ribosome abundance increases translational noise. In our simulation, increasing translational noise or reducing translation rates results in increasing the cycle-to-cycle variability in circadian period. This aligns with previous stochastic modeling, which found that as the average number of molecules in the cell is increased and interactions become more frequent, oscillator period variability decreased^81^. These results mirror the circadian variability observed between and within AD patients. Disrupted circadian timekeeping could stem from weakened molecular oscillators, reduced synchronization among pacemakers, or impaired environmental entrainment. Our findings suggest that dysregulated output rhythms in AD patients may partly result from altered ribosomal dynamics.

Overall, this study highlights a three-pronged interaction that may exacerbate AD progression (Fig. 7F): (1) AD pathology, particularly amyloid plaque formation, leads to widespread ribosomal dysregulation and reduced protein translation capacity; (2) sleep and circadian disruption, early AD symptoms, further amplify this dysfunction; and (3) altered translation feeds back on the circadian clock, increasing timekeeping variability.

While it has been established that molecular circadian disruption through BMAL1 KO worsens pathology in AD model mice, the influence of behavioral circadian disruption on AD pathology is yet to be established.

Several limitations should be considered in evaluating this work. Our data were ordered informatically to address challenges of real-world data and the biology of AD: (1) Time of death (TOD) is often unreported in large-scale studies and may be inaccurately estimated in home environments. (2) AD patients’ internal circadian phases are often poorly coupled to external time. The phase reported by CYCLOPS reflect a samples relative position in the circadian molecular cycle rather than its phase with respect to an external timing cue. As such our work complements, but is distinct from studies tied to external timing cues which suggest that AD affected subjects are, on average, phased delayed-relative to CTLs^68^. Moreover, our analysis cannot distinguish rhythms directly driven by the molecular oscillator from those imparted by daily behaviors or environmental cycles. This restricts our ability to distinguish circadian rhythms from broader diurnal oscillations. While informatic reconstructions require cautious interpretation, we employed multiple strategies to mitigate noise and validate our results. Our CYCLOPS ordering was based on a specific population of excitatory neurons with a strong circadian signal and a curated gene list known to cycle in the human and mouse cortex. CYCLOPS 2.0 controlled for sequencing batch and disease state in the ordering process. We also adjusted for post-mortem interval, sex, and sequencing batch in the identification and downstream analysis of rhythmic molecules. Additionally, an alternative algorithm, CHIRAL, yielded similar sample phase assignments. Perhaps most importantly, our analysis reproduced the established, relative acrophases of core clock transcripts and aligned with measured timing of clock output genes in human cortex.

Our pipeline used the sample phases estimated from excitatory neurons for the analysis of other cell types. This strategy was informed by mouse data showing that, in both WT and AD models, core clock transcripts are synchronized across cell types. If cell-type clocks had fixed phase offsets (e.g., BMAL1 peaking at different times in neurons vs. glia), we would have detected this pattern. If neuron and glial clocks were entirely unsynchronized, an ordering based on neuronal time would very likely fail to reconstruct appropriate core clock dynamics in other cells.

Our human samples were obtained post-mortem. While ROS/MAP has worked to optimize collection procedures, prolonged post-mortem intervals and other perimortem factors can affect RNA quality and lead to unique molecular differences before death. However, these should be shared between both the AD dementia and CTL groups and we included post-mortem interval as a confounding variable in our regression analyses.

For follow-up studies, we used the plaque-bearing APP^swe^/PS1^ΔE9^ model, as our informatic analysis of human data highlighted a link between ribosomal dysregulation and plaque burden. However, our results may be specific to this model and age group. Previous studies have shown that CJL protocols can influence both sleep duration and circadian rhythmicity, making it difficult to completely distinguish the effects of these distinct but related factors. The use of polysome profiling provided valuable insights into the functional consequences of the APP^swe^/PS1^ΔE9^ genotype and the CJL protocol on protein translation, but this approach has inherent limitations. While polysome analysis offers a broad measure of translational engagement, it does not resolve changes at the level of specific proteins. Moreover, polysome profiling does not distinguish between stalled and actively translating ribosomes.

As with all informatic analyses, correlation does not imply causation. We do not propose that altered ribosomal biogenesis or oxidative phosphorylation rhythms are the causative insults underlying AD. Much of the transcriptomic signature we observe may reflect downstream consequences of AD pathology. Instead, we acknowledge that sleep and circadian disruption—consequences of AD—may partly drive these molecular changes. Nonetheless, addressing these downstream effects could help mitigate AD-related decline. Perhaps more importantly, follow-up mouse experiments confirmed that the plaque producing APP/PS1 genotype causally alters ribosomal biogenesis and oxidative phosphorylation rhythms. The diminished cycling and reduced basal abundance of ribosomal transcripts observed in AD patients were mirrored in AD model mice. We observed a reduction in the abundance of ribosomal components at both the transcript and protein level. Our polysome analysis suggests that this functionally impacts protein translation. Unlike human AD, APP/PS1 mice exhibit minimal sleep and circadian disruption under controlled conditions^16^. However, a chronic jet lag protocol that combines elements of sleep deprivation and circadian disruption exacerbated AD-related declines. These human and mouse data suggest stabilizing circadian rhythms could have therapeutic potential. However, without randomized clinical studies, the impact of circadian or sleep-wake interventions on AD patients remains uncertain.

**Table 1.** Summary statistics of included ROSMAP subjects.

| Condition | CTL | AD | Total |
| --- | --- | --- | --- |
| Sample size (n) | 250 | 159 | 409 |
| Mean age at death (yrs) | 86.4 | 88.0 | 87.0 |
| Standard dev. age at death (yrs) | 4.8 | 3.7 | 4.5 |
| Percent male (%) | 34.8 | 28.3 | 32.3 |
| Median PMI (hrs) | 6.0 | 5.8 | 6.0 |

## Resource Availability

The dataset(s) supporting the conclusions of this article are available in the AD Knowledge Portal repository https://www.synapse.org/#!Synapse:syn3219045, and https://www.synapse.org/Synapse:syn26007830. Access is restricted to preserve anonymity and requires a data use certificate (DUC) to be submitted https://adknowledgeportal.synapse.org/Data%20Access.

## Acknowledgements.

This work was supported by NIA R01 AG068577 (RCA) and NIA R01AG054517 (ESM), In addition, AS was funded in part by the BrightFocus Foundation grant number A2024031F.

ROSMAP (DAB) is supported by P30AG10161, P30AG72975, R01AG15819, R01AG17917, U01AG46152, and U01AG61356.

The results published here are in whole or in part based on data obtained from the AD Knowledge Portal Community Data Contribution Program.

ROSMAP resources can be requested at https://www.radc.rush.edu and www.synpase.org.

Metabolomics data generated by the Alzheimer’s Disease Metabolomics Consortium (ADMC; ADMC members list https://sites.duke.edu/adnimetab/team/), led by Dr. Rima Kaddurah-Daouk at Duke University, using biospecimens provided by the Rush Alzheimer’s Disease Center, Rush University Medical Center, Chicago.

## Author Contributions

Conceptualization: RCA, ESM, VM, HCH, and AS.

Data & Data Curation: HCH, PWS, VM, DAB

Investigation: HCH, AS, PWS, LBM, and JDW

Formal Analysis: HCH, AS, ESA, and RCA

Software: HCH, JAH, RCA

Writing – Original Draft: HCH, AS, ESA, and RCA

Writing – Review & Editing: All authors

## Materials and Methods

### Animals

All animal procedures were approved by the Institutional Animal Care and Use Committee (IACUC) at Washington University and conducted in accordance with the guidelines of the Association for Assessment and Accreditation of Laboratory Animal Care (AALAC). Male B6;C3-Tg(APPswe,PSEN1dE9)85Dbo/Mmjax hemizygous mice (PSAPP, Stock #004462) and female B6C3F1/J mice (Stock #100010) were obtained from Jackson Laboratory (Bar Harbor, ME) and bred in-house at Washington University in St. Louis. Genotypes were confirmed through tail biopsies, and the study included both male and female PSAPP carriers and non-carriers. The PSAPP mouse model replicates key aspects of Alzheimer’s disease by incorporating humanized sequences encoding amyloid precursor protein (Mo/HuAPP695swe) with Swedish mutations (K595N/M596L) and a mutant form of human presenilin 1 (PSEN1dE9), which features an exon-9 deletion^84,85^. Mice were group housed and aged to 12 months before being transferred to specialized rooms for simulated chronic jetlag (CJL) induction. They were acclimated for one week in well-ventilated, soundproof, and light-tight circadian cabinets (Actimetrics, Lafayette, IN) with computer-controlled light–dark cycles. The control groups followed a standard light-dark (LD) schedule with light exposure from 0600 hrs to 1800 hrs, while the CJL group was subjected to a 6-hour phase advance each week for three months (12 cycles) (Fig 6A). Following CJL exposure, mice were allowed a one-week re-entrainment period before tissue collection. Perfusions were performed between Zeitgeber Time (ZT) 5 and ZT7 (1100 to 1300 hrs), during which mice were deeply anesthetized with intraperitoneal pentobarbital (150 mg/kg) and perfused with ice-cold Dulbecco’s modified phosphate-buffered saline (DPBS) containing 3 g/L heparin.

The left hemisphere of each brain was immersion-fixed in 4% paraformaldehyde for 24 hours at 4°C, followed by cryoprotection in 30% sucrose in PBS for 24 hours at 4°C. Fixed brains were sectioned into 40 µm coronal slices using a freezing sliding microtome (SM1020R; Leica) and stored in cryoprotectant solution (30% ethylene glycol, 15% sucrose, and 15% phosphate buffer in distilled water). For the molecular and biochemical analyses, cortices from the right hemispheres were dissected, flash-frozen, and stored at –80°C.

### snRNA-seq Data

Human data came from participants in the Religious Orders Study or Rush Memory and Aging Project (ROSMAP)^38^. All ROSMAP participants enrolled without known dementia and agreed to detailed clinical evaluation and brain donation at death. Both studies were approved by an Institutional Review Board of Rush University Medical Center.

Each participant signed informed and repository consents and an Anatomic Gift Act. We analyzed single-nucleus RNA sequencing data including 250 aged subjects without dementia (CTL) and 159 with AD dementia. Subject condition labels were based on the clinical consensus diagnosis of cognitive status at time of death (COGDX), provided by ROSMAP. Specifically, CTL subjects were defined by COGDX < 3 and AD subjects were defined by 3 < COGDX < 6 (COGDX of 6 were excluded for having dementia from a source primarily other than AD). Of the twelve subjects that had duplicate sequencing batches, we used sequencing profiles from the batch with more counts. Sequencing, processing reads into counts, and cell type labelling were made available from previous work^86^. Single-nucleus RNA-seq reads and processed counts, with cell-type classifications, are available at Synapse (<u>syn31512863</u>).

### Quantifying Circadian Signal with CCD

The clock correlation distance (CCD)^40^ is a marker of the core clock circadian signal in unordered data. CCD quantifies the circadian rhythm “choreography” in a population by comparing the gene-gene correlation matrix of 12 core clock genes with an established reference correlation matrix. Given the upper triangle of the correlation matrix from a population, C, and the upper triangle of the established reference correlation matrix, R, CCD is calculated as follows:

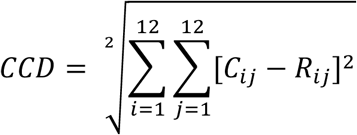

We systematically calculated the CCD of different combinations of excitatory neuron subclusters from the CTL subjects to identify specific subclusters with the strongest circadian signal. Specifically, we calculated the CCD for different combinations of the sixteen subclusters, considering group sizes k = 1,2,3,4,5 (,^$&^- combinations for each k). The combined population of excitatory neurons 3-5 produced the correlation matrix most close to the reference (CCD = 3.26).

### Aggregating and Normalizing Counts

Single-nucleus RNA counts were aggregated with the Seurat package^87^ by cell type and subject to create cell type specific pseudo bulk data. Mitochondrial genes were excluded. Lowly expressed genes were filtered out with the filterByExpression function (default parameters) in edgeR^88–91^. Trimmed Mean of M-values (TMM) were obtained using calcNormFactors() method in edgeR.

### Differential Expression

For differential expression analysis, sex, PMI, and sequencing batch were included as covariates. The two sequencing batches were composed of those sequenced by the Broad Institute’s Genomics Platform, and those resequenced at the New York Genome Center, as described by Green et al^39^. Standard edgeR differential expression was performed between our CTL and AD subjects for each cell type. First we estimated the dispersions using the estimateDisp() function. We fit negative binomial generalized linear models with the glmQLFit() function. Finally, differential expression was determined with the quasi-liklihood (QL) F-test using the glmQLFTest() function.

### Determining Subject Phases

We inferred the internal circadian phases of subjects using the CYCLOPS 2.0 algorithm. We applied CYCLOPS 2.0 to cell type-specific pseudo-bulk data from combined excitatory neuron subclusters 3-5. We excluded 7 subjects (4 CTL, 3 AD) with fewer than 10 excitatory subcluster 3-5 cells total from the ordering process.

A set of genes likely to cycle in a dataset is required for the successful application of CYCLOPS 2.0. We synthesized such a set by taking the union of three sources: Cycling genes from WT mice bulk tissue^37^ (JTK BH.q < 0.01), genes that cycle in 10 of 12 tissues in the mouse atlas^4^, and the top 25 genes cycling in human BA11^33^ (ranked by cycling p value).

Subject diagnosis, AD dementia or CTL, and sequencing batch were provided as covariates to CYCLOPS 2.0. The two sequencing batches were composed of those sequenced by the Broad Institute’s Genomics Platform, and those resequenced at the New York Genome Center, as described by Green et al^39^. CTL samples were ordered in conjunction with the AD samples. The CTL and AD samples were projected onto the CTL samples’ eigengene space to emphasize the circadian variation in AD samples, the same approach used in Anafi et al. 2017^35^.

The CYCLOPS parameters were left at default values. Performance metrics of CYCLOPS were assessed and found to be adequate (Stat_Err_ = 0.20 with p value of 0.02, and smoothness 0.84). The Stat_Err_ metric was calculated on a CYCLOPS model trained with just CTL samples, whose phase predictions generally matched the CYCLOPS model jointly trained with AD and CTL samples.

We also inferred circadian phases of these subjects using CHIRAL, an alternative to CYCLOPS. We provided the same cycling genes to CHIRAL as we did to CYCLOPS. The two orderings were well correlated (JR circular cor = 0.84, Fig S2D).

### Metabolomics and MetaboAnalyst

Metabolomics data were generated by the Alzheimer’s Disease Metabolomics Consortium (ADMC members list https://sites.duke.edu/adnimetab/team/), led by Dr. Rima Kaddurah-Daouk^92–94^. Metabolite data was available for 204/409 of the individuals we studied (118 CTL and 86 AD dementia subjects). Previously processed data is available on Synapse (syn26007830). Unlike the snRNA-seq data, the metabolite data is not specific to cell type. We assigned the CYCLOPS predicted phases to these metabolic samples and performed the same cycling/differential cycling analysis as before. The MetaboAnalyst Joint Pathway Analysis tool was used to infer metabolic pathways associated with cycling, differentially cycling, or differential MESOR transcripts and metabolites in CTL and AD. The transcripts we provided to this tool were from the full pseudo-bulk transcriptome, combining counts from all cell types. For the analysis of joint pathways cycling in CTL (Fig S6E), we used transcripts and metabolites cycling with BH.q < 0.1. In our analysis of differential MESOR pathways (Fig S6G), we used transcripts and metabolites cycling with BH.q < 0.1. For the analysis of joint pathways differentially cycling (Fig S6F), we used transcripts differentially cycling in the full pseudo-bulk transcriptome with BH.q < 0.3 and metabolites differentially cycling with uncorrected p-value < 0.05. All settings on the Joint Pathway Analysis tool were left at default.

### Pathway Analysis with Enrichr and fGSEA

To get a more complete understanding of the biological pathways associated with a list of transcripts we used both the Enrichr^42–44^ API and the fGSEA R package^41^.

The Enrichr platform uses and overrepresentation approach. We used the KEGG_2021_Human Enrichr library. The Enrichr API allows users to provide a background set of genes. We used all transcripts expressed in a cell type as the Enrichr background when testing for pathways associated with cycling genes. We used all the genes tested for differential rhythms in a cell type (i.e. genes with cycling BH.q < 0.1 and amplitude / MESOR ≥ 0.2 in AD or CTL) as the background when we tested for pathways associated with differentially rhythmic genes.

fGSEA uses an enrichment approach to pathway analysis. We called the fgsea function with arguments nPermSimple =10,000, gseaParam =1, minSize = 10, maxSize = 500. We used the MsigDB gene matrix transpose files (gmt files) for Hallmark (h.all.v2024.1.Hs.symbols.gmt) and KEGG (c2.cp.kegg_legacy.v2024.1.Hs.symbols.gmt) libraries. The code for using Enrichr and fGSEA can be found on our Github repository.

### Immunofluorescence (IF) and imaging

Immunofluorescence staining was conducted on two brain sections per mouse. Sections were blocked in 3% donkey serum for 60 minutes and incubated overnight at 4°C with the RPS6 primary antibody. The next day, sections were counterstained with a fluorescent secondary antibody and DAPI. Image acquisition was performed using a Nikon AXR with NSPARC Confocal Microscope and NIS Elements software (v5.4, Nikon Instruments Inc.). Images were captured in a sequential mode using 0.3 µm z-steps, maintaining uniform settings for pinhole size, laser power, and detector configurations. Quantification of fluorescence was carried out using Imaris 10.1 software (Oxford Instruments). Brain regions of interest included the dentate gyrus and isocortex.

### Protein extraction and immunoblotting

Frozen cortical tissue was homogenized in ice-cold radioimmunoprecipitation assay buffer supplemented with a 1X Halt Protease Inhibitor Cocktail. The soluble fractions were collected, aliquoted, flash-frozen, and stored at −80°C. Protein concentration was determined using a bicinchoninic acid (BCA) Protein Assay Kit (Thermo Scientific). Equal amounts (10 μg) of total protein were mixed with NuPAGE sample reducing agent (Invitrogen), combined with 4X Laemmli sample buffer, and separated on Novex 16% Tricine gels (Invitrogen) in Tricine SDS running buffer supplemented with NuPAGE Antioxidant. Proteins were transferred to polyvinylidene fluoride membranes, which were blocked in 5% milk in Tris-buffered saline with 0.1% Tween-20 (TBST) for 60 minutes at room temperature. Primary antibodies were used at a 1:1000 dilution, targeting RPS6, RPL7A, RPS27A, RPL10A, and beta-tubulin. Membranes were incubated with primary antibodies overnight at 4°C, followed by washing and incubation with horseradish peroxidase (HRP)-conjugated secondary antibodies for 1 hour at room temperature. Protein detection was performed using Pierce ECL Western Blotting Substrate or Lumigen ECL Ultra, and imaging was carried out on a ChemiDoc Imaging System (Bio-Rad).

For sequential blotting of multiple proteins, membranes were stripped using Restore PLUS Western Blot Stripping Buffer (Thermo Scientific) at 37°C for 30 minutes with agitation. After stripping, membranes were washed with deionized water, followed by three washes in TBST (10 minutes each) before blocking and reprobing.

### m47S RT-PCR

Quantitative PCR for 47S pre-rRNA transcripts was conducted following the method described by Cui and Tseng^95^ with modifications for murine 47S pre-rRNA sequences. Total RNA was extracted from frozen cortical tissue using Trizol and the PureLink RNA Mini Kit (Invitrogen) per the manufacturer’s instructions. First-strand cDNA synthesis was performed using the SuperScript III First-Strand Synthesis System (Invitrogen) with an rDNA-specific primer (5′-CGTGGCATGAACACTTGG-3′). Real-time PCR was performed using PowerUp SYBR Green Master Mix (Applied Biosystems) on a QuantStudio 3 system (Applied Biosystems). The m47S pre-rRNA copy number was measured using the following mouse-specific primers:

- Forward: 5′-CTGACACGCTGTCCTTTCCC-3′
- Reverse: 5′-GTGAGCCGAAATAAGGTGGC-3′

A standard curve was generated from serial dilutions of a known quantity of subcloned cDNA to quantify transcript copy numbers.

### Polysome Fractionation

Polysome fractionation was performed as described by Strezoska et al^61^. Briefly, 25–45 mg of frozen cortical tissue was lysed in 250 µL of polysome lysis buffer (20 mM Tris, pH 7.2; 130 mM KCl; 10 mM MgCl₂; 2.5 mM DTT; 0.5% NP-40; 0.5% sodium deoxycholate; 0.2 mg/mL heparin; 10 µg/mL cycloheximide; 200 U/mL RNasin). The tissue was incubated on ice for 5 minutes, dissociated via serial needle aspirations (ten times through an 18-gauge needle, followed by five times through a 25-gauge needle), and incubated on ice for an additional 15 minutes. Lysates were cleared by centrifugation at 8000×g for 10 minutes at 4°C. Cleared lysates were layered onto a 12 mL 17–47% (w/w) sucrose gradient prepared in sucrose base buffer (10 mM Tris, pH 7.2; 60 mM KCl; 10 mM MgCl₂; 1 mM DTT; 0.1 mg/mL heparin) and centrifuged at 36,000 rpm for 3 hours at 4°C in an SW41 Ti swinging-bucket rotor.

Gradient fractionation was performed using a Biocomp Instruments Gradient Fractionation System (Tatamagouche, NS, Canada), with UV absorbance at 260 nm recorded using a Triax Flow Cell and associated software (Biocomp Instruments). Polysome traces were aligned and quantified using the QuAPPRo tool^62^.

### Mathematical Model of Mammalian Clock

We adapted a previously published mathematical model of the mammalian core clock oscillator^63^. This model includes seven molecular species: Bmal1 mRNA, BMAL1 protein, nuclear BMAL1, active BMAL1 (BMAL1-CLOCK complex), Per2/Cry mRNA, PER2/CRY protein complex, and nuclear PER2/CRY complex.

We modified this deterministic model in two ways. First, to simulate translational slowing, we introduced t_slow_, a parameter that scales the translation rates of BMAL1 and PER2/CRY proteins without replacing the original translation terms. Second, to incorporate stochasticity in translation, we added a diffusion term driven by a Wiener process, t_noise_, introducing Gaussian noise in the translation rate of BMAL1 and PER2/CRY proteins. To determine a reasonable choice for this noise parameter in control samples, we used empirical data: the standard deviation of the circadian period in a control human population is approximately 0.2 hours^96,97^. We found that setting t_slow_ = 0.03 reproduced this variability.

We simulated 20 individuals for each choice of t_slow_ and t_noise_ parameters, throwing away the first 20 oscillations in each simulation to let the models stabilize. Individual cycle periods were identified based on the extrema of Bmal1 mRNA, which remained unaffected by changes to t_slow_ and t_noise_. The modified oscillator model was implemented in Julia, using the DifferentialEquations package and the code is available on our GitHub.

### Rhythmicity, Differential Rhythmicity, Differential MESOR

Transcripts from CTL and AD samples were tested for rhythmicity independently. Transcripts with a BH.q < 0.1 and an amplitude ratio (amplitude / MESOR) ≥ 0.2 were considered rhythmic. Transcripts that were rhythmic in either CTL or AD were tested for differential rhythmicity. Transcripts with a differential rhythmicity BH.q value < 0.3 were considered differentially rhythmic. All transcripts were tested for differential MESOR.

Cosinor regression is a common technique for evaluating the fit of data to a sinusoidal waveform. Using this technique, we took the predicted subject phases from CYCLOPS, 𝜃 (0 ≤ 𝜃 ≤ 2𝜋), and obtained a wave of best fit for individual transcripts. Importantly, we test if the data is better fit by a waveform than a straight line. Given subject phases from CYCLOPS, 𝜃, we fit the abundance of a molecular species, Y, with two nested, linear mixed models (Eqs.1-2). Both models include sequencing batch, sex, and PMI as covariates. The null model predicts expression of a species, Y, as a function of the sequencing batch variable, 𝑏, a continuous PMI variable, 𝑝, a discrete sex variable, 𝑠, and an intercept, 𝛽_(_. The expanded model has additional sinusoidal functions of CYCLOPS phase. The idea is to determine if the data ordered by CYCLOPS is better fit with a sinusoid than without, assessed by the F test. This is repeated for each transcript independently. The null hypothesis of the F test is that sinusoid coefficients 𝛽_)_= 𝛽_*_ = 0.

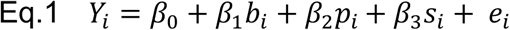

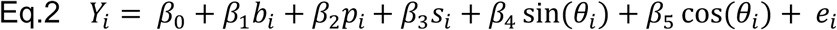

Assuming a gene is significantly better fit by Eq.2, the amplitude, A, of the rhythm can be found:

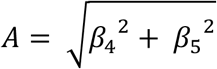

The acrophases of the rhythm, 𝜙, can be found using the trigonometric identity:

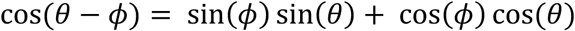

so

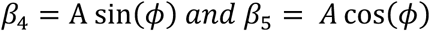

Therefore, the acrophases of the rhythm, 𝜙, is:

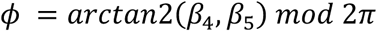

After the cycling genes are separately identified in AD and CTL subjects, we assess differential rhythmicity of transcripts with nested linear models. The F test here has the null hypothesis that 𝛽_7_= 𝛽_8_ = 0.

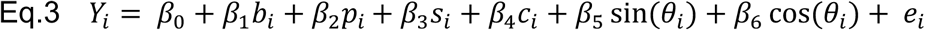

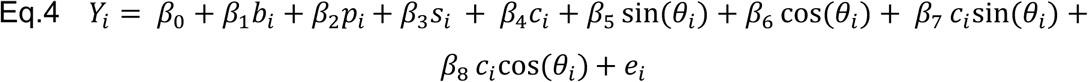

Where 𝑐_i_ is the sample condition (CTL or AD) for sample 𝑖. Only genes that were cycling in CTL or AD are tested for differential rhythmicity (BH.q < 0.1 and amplitude / MESOR ≥ 0.2 in AD or CTL).

We also tested differential rhythms stratifying samples by Braak score and CERAD score, pathology measures provided by ROSMAP. For this we used the same differential rhythms equations above (Eq.3-4), but replaced the sample condition, 𝑐_i_, with a binned Braak score (0, I, II), (III, IV), (V, VI) or a binned CERAD score (1,2), (3, 4).

Differential MESOR analysis was also performed. This tests for differences in a transcript’s MESOR between CTL and AD. One can think of this as a circadian-corrected differential expression analysis. This too is done with two nested linear models and an F test. The F test here has the null hypothesis that 𝛽_6_ = 0.

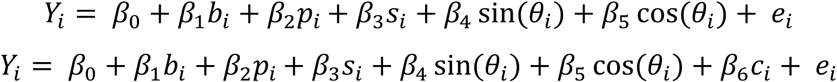

### Kendall rank correlation coefficient

When quantifying Western blotting, imaging, and polysome profiling, we used the Kendall rank correlation coefficient to understand the relationship between ordinal experimental group and the dependent variable. Like standard Pearson correlation, Kendall’s tau measures the strength and direction of association between two variables. However, unlike Pearson correlation, Kendall’s tau is nonparametric and does not assume a linear relationship, making it particularly suited to our analysis. Because the experimental groups (WT+LD, APP+LD, APP+CJL) represent ordinal categories, we rank them as follows: WT+LD < APP+LD < APP+CJL.

Kendall’s tau is preferable to other rank-based correlation metrics, such as Spearman’s correlation coefficient, because it is better equipped to handle ties in ranks. Additionally, Kendall’s tau provides a p-value for the association, which we indicate on our plots using slanted lines to denote statistical significance.

---

**Figure S1.**
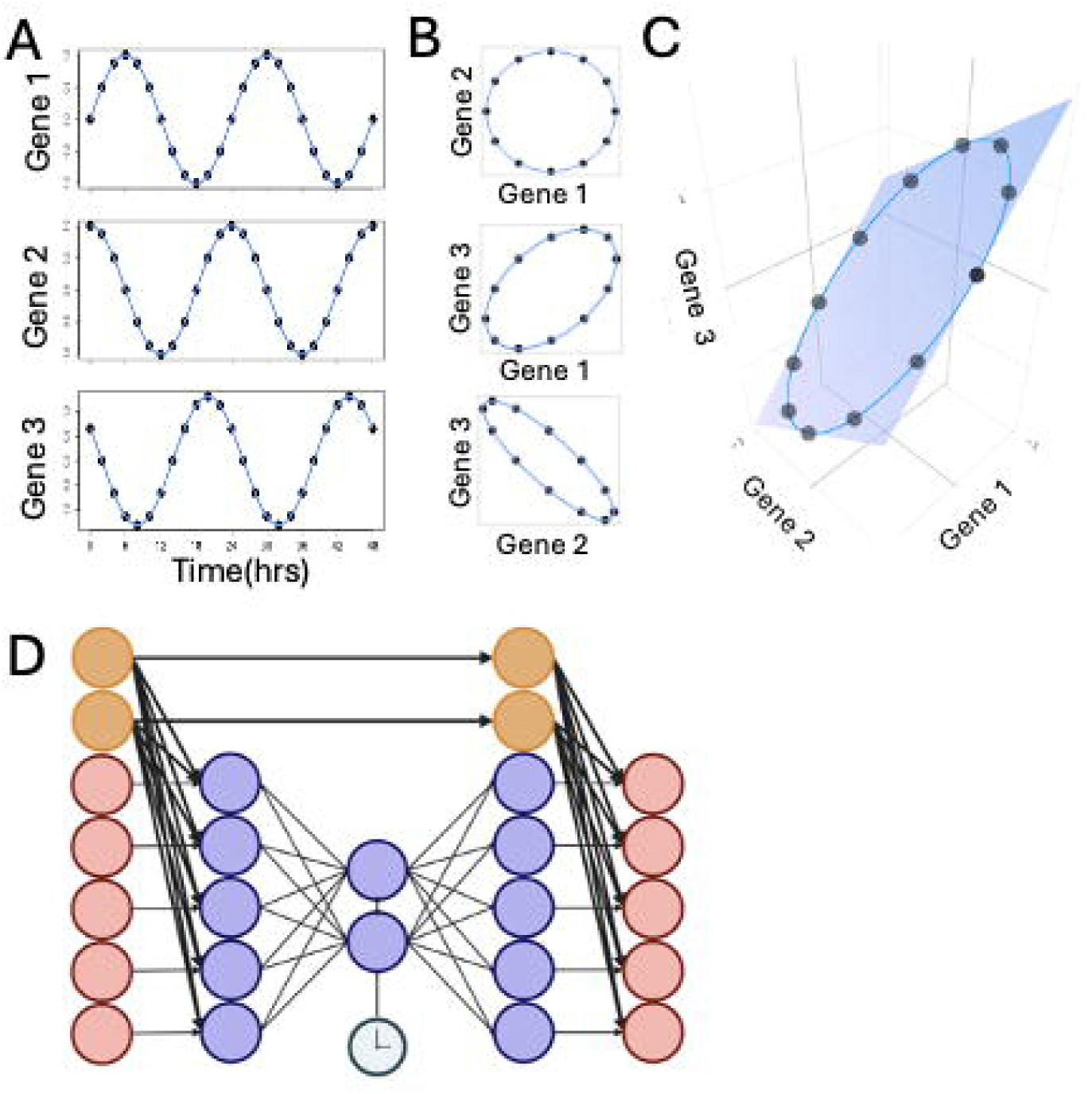
Principles of the CYCLOPS algorithm, related to Figure 1. **(A)** Three rhythmic, out of phase genes as a function of time. **(B)** pairwise comparisons of rhythmic out of phase genes form ellipses. **(C)** Plotting the expression of three cycling genes against each other, a 2-dimensional ellipse again arises. The expression of hundreds or thousands of circadian genes lies on a single ellipse in gene expression space. The relative position of each sample on the ellipse in gene space represents its relative position in the circadian cycle. **(D)** CYCLOPS uses an autoencoder to identify circular structure in high-dimensional data. CYCLOPS 2.0 allows the autoencoder to adjust these data for non-circadian confounders such as sequencing batch and disease state.

**Figure S2.**
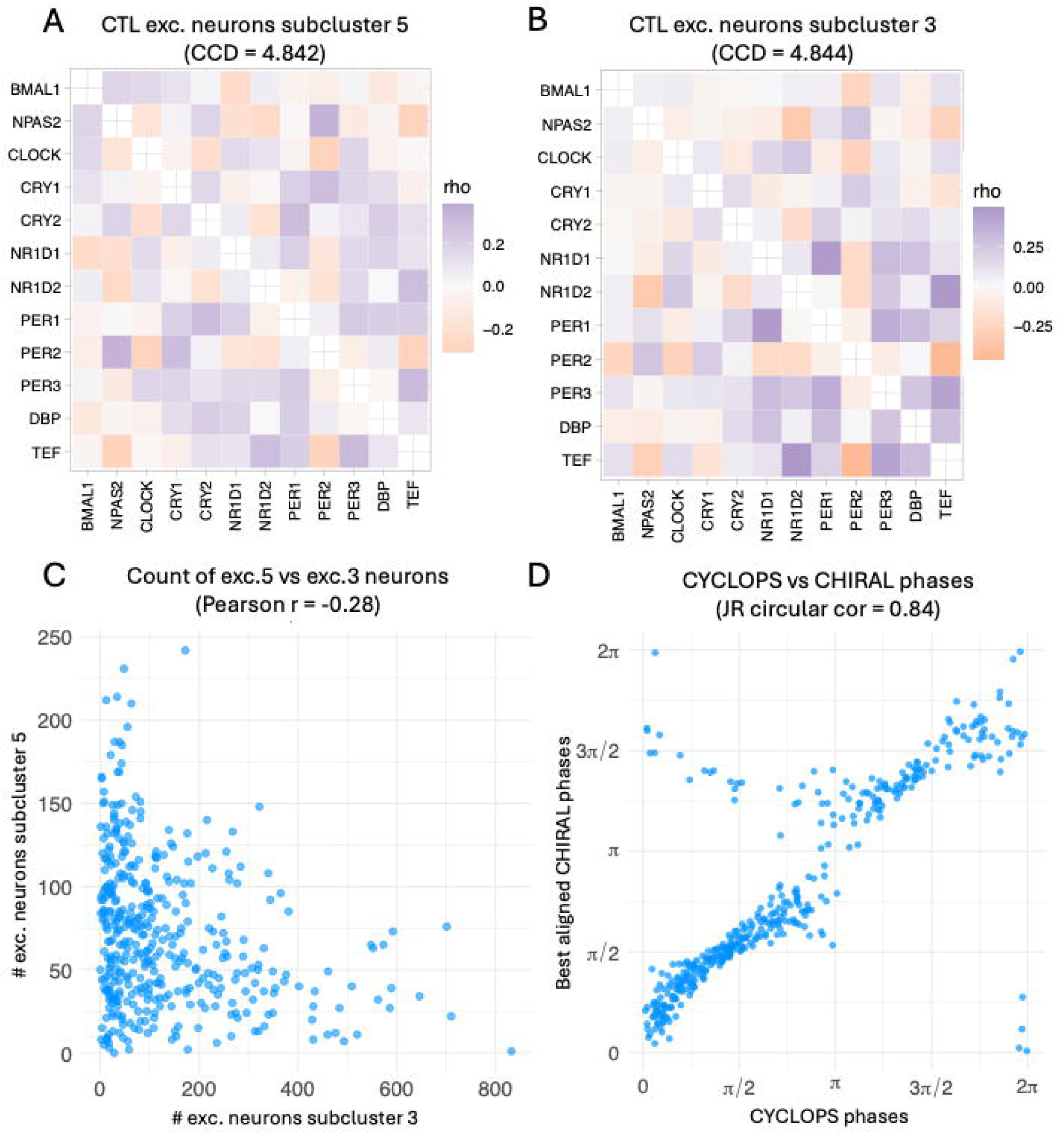
Selection of cell subclusters for ordering and comparison with alternative ordering algorithm, related to Figure 2. **(A, B)** Gene-gene correlation matrices for excitatory neuron subcluster 5 (CCD = 4.84) and excitatory neuron subcluster 3 (CCD = 4.84), respectively. The matrices show that neither subcluster individually resembles the established reference matrix, as indicated by their higher CCD values compared to the combined excitatory neuron subtypes 3-5 (CCD = 3.26). **(C)** Counts of excitatory subcluster 5 neurons versus excitatory subcluster 3 neurons for each individual. A negative correlation is observed between the counts of the two subclusters (Pearson r = −0.28). **(D)** CYCLOPS-predicted phases compared to CHIRAL-predicted phases for all subjects (CTL and AD). CYCLOPS and CHIRAL phases are in strong agreement (Jammalamadka ranked circular correlation = 0.84). Horizontal axis: CYCLOPS-predicted phases. Vertical axis: CHIRAL-predicted phases, aligned to CYCLOPS phases.

**Figure S3.**
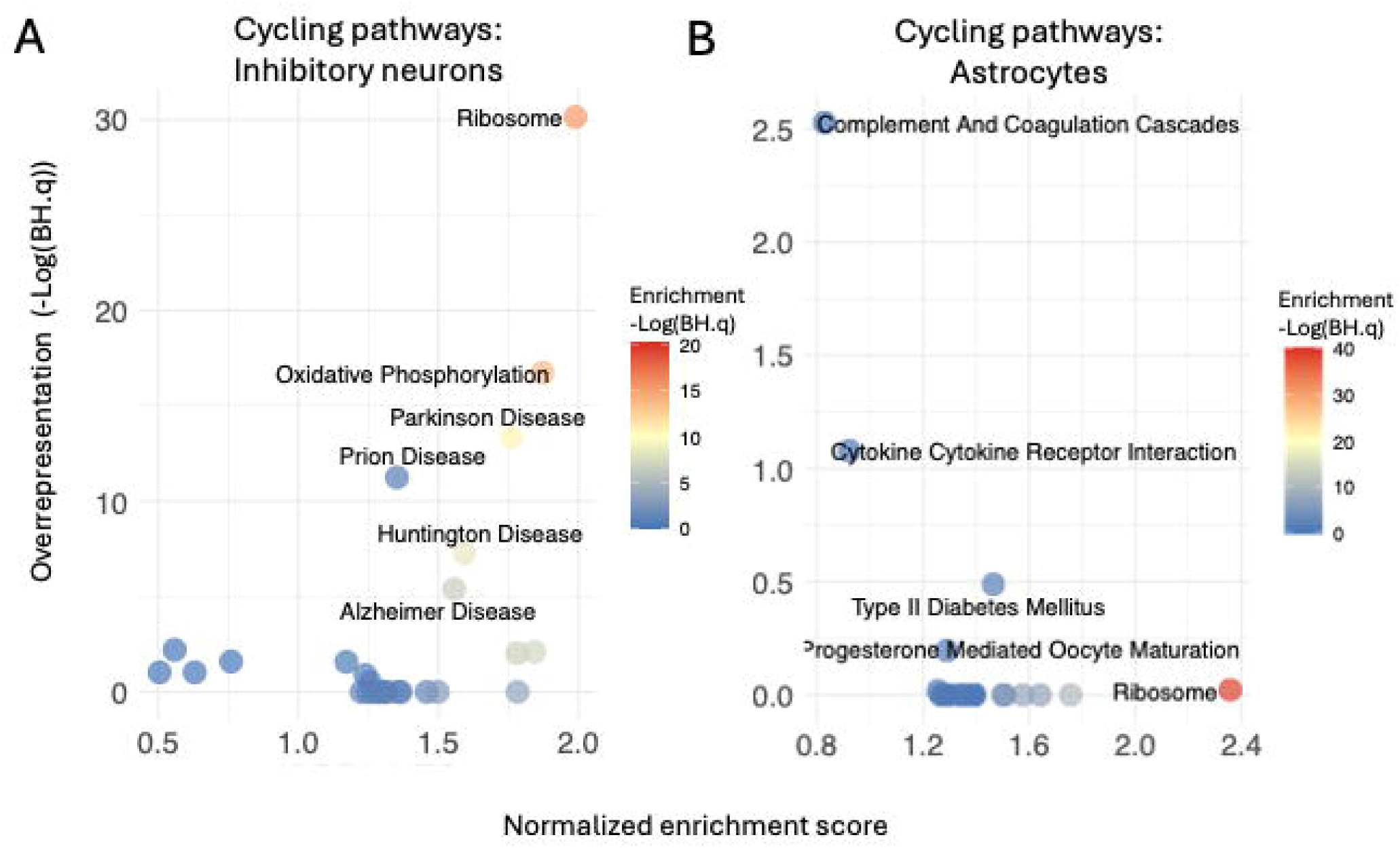
Pathway-level cycling analysis in CTL samples across cell type, related to Figure 2. **(A)** Inhibitory neurons, **(B)** astrocytes. Two complementary approaches were applied: enrichment analysis using fGSEA and overrepresentation analysis using Enrichr. The horizontal axes show the fGSEA normalized enrichment score, while the vertical axes represent the significance of overrepresentation (-Log(BH.q)) derived from Enrichr. The color scale indicates the significance of enrichment (-Log(BH.q)) as calculated by fGSEA.

**Figure S4.**
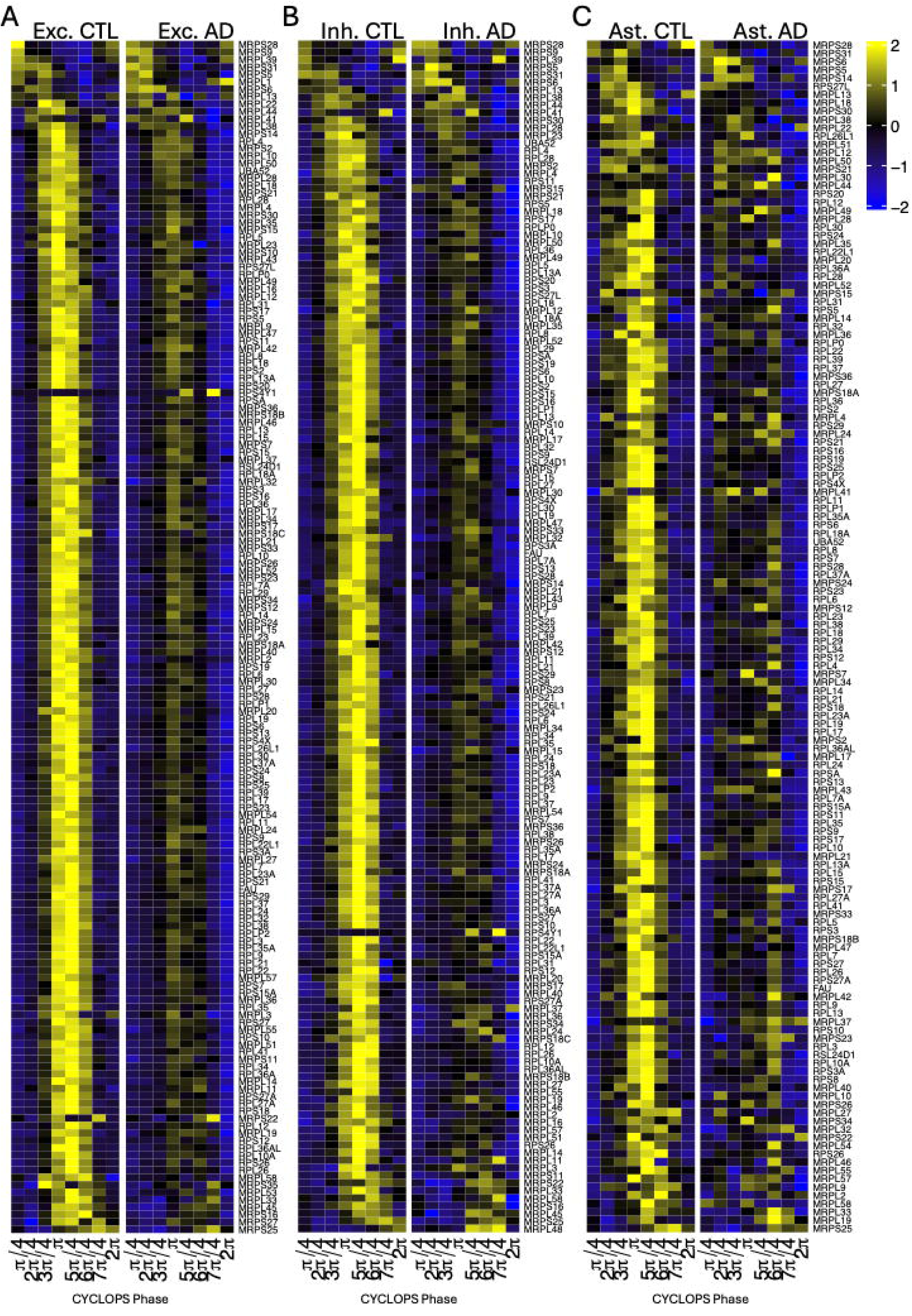
Circadian heatmaps of ribosomal genes in human cell types, related to Figure 5. **(A)** Excitatory neurons, **(B)** inhibitory neurons, and **(C)** astrocytes. Heatmaps show cycling ribosome genes in CTL (left) and AD (right) individuals. Rows are ordered by CYCLOPS-predicted acrophases, and columns represent binned CYCLOPS phases.

**Figure S5.**
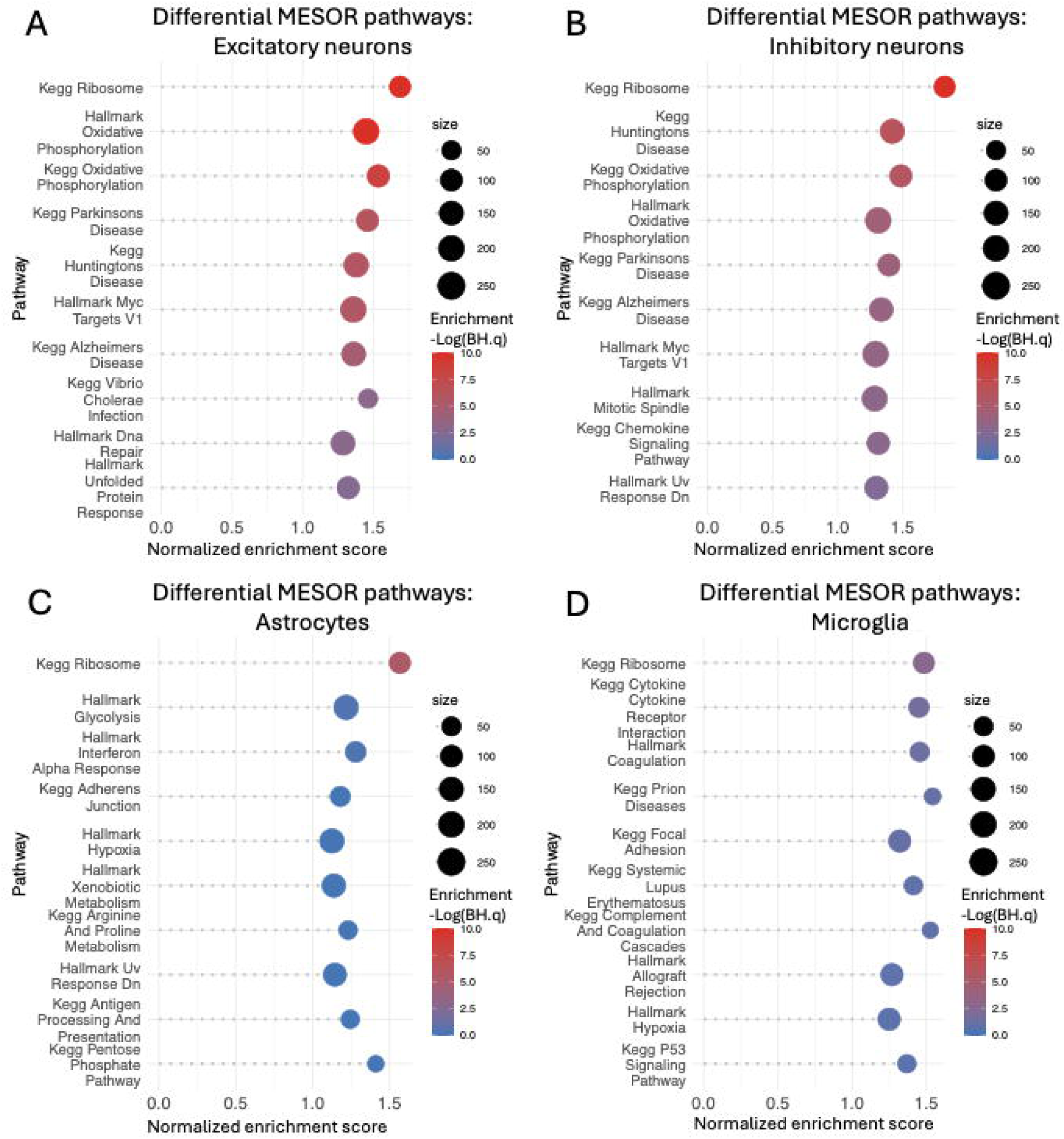
Pathway analysis of differential MESOR transcripts across cell type, related to Figure 5. **(A)** Excitatory neurons, **(B)** inhibitory neurons, **(C)** astrocytes, and **(D)** microglia. Vertical axis: KEGG and Hallmark pathways. Horizontal axis: fGSEA normalized enrichment score. Dot size represents number of genes within a pathway, and color represents enrichment significance.

**Figure S6.**
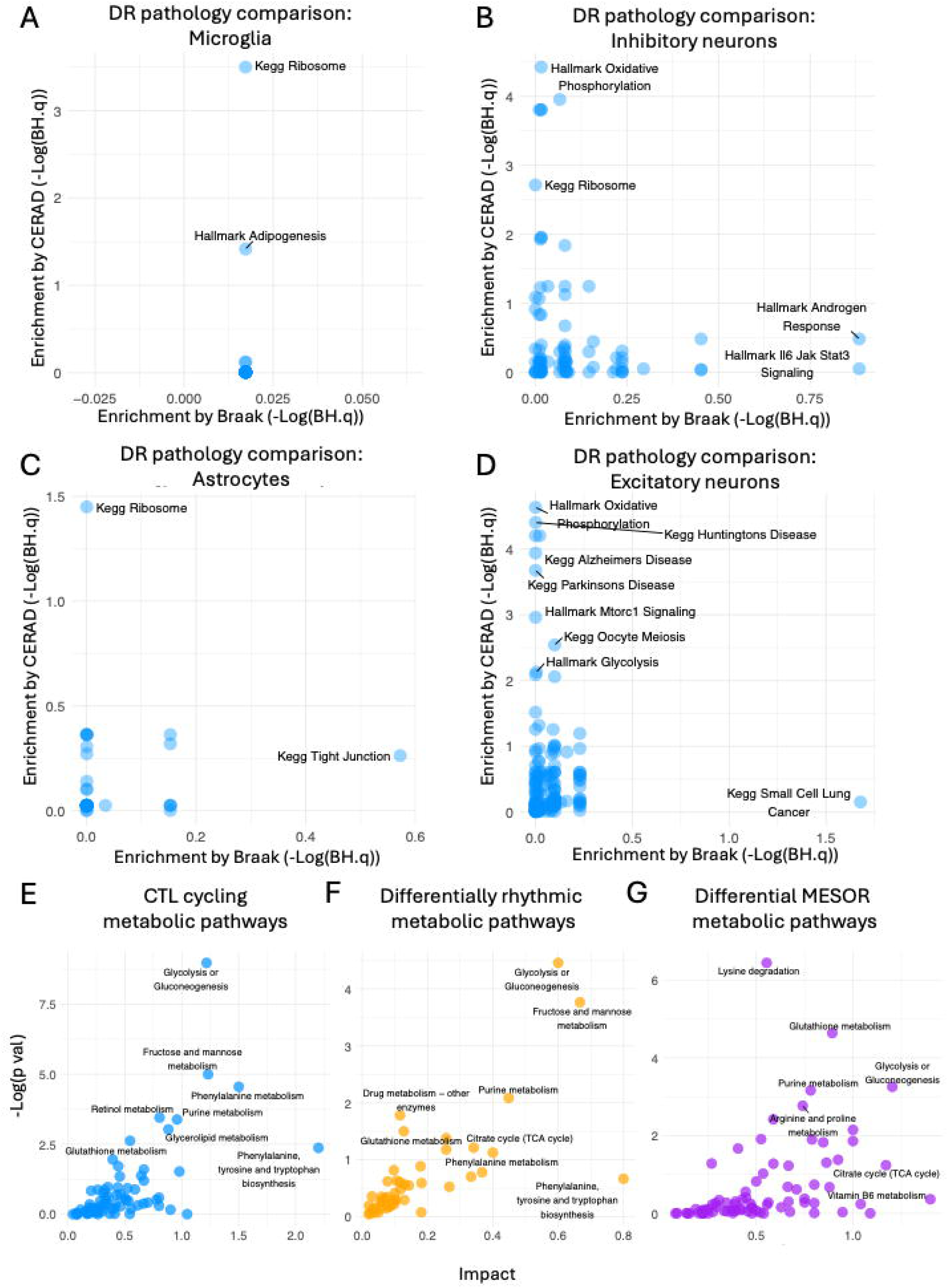
Differentially rhythmic transcriptional and metabolic pathways in different cell types and pathologies, related to Figure 5. **(A-D)** Pathway analysis of differentially rhythmic genes stratified by CERAD and Braak scores for **(A)** microglia, **(B)** inhibitory neurons, **(C)** astrocytes, and **(D)** excitatory neurons. The vertical axes represent the fGSEA enrichment significance (-Log(BH.q)) of genes differentially rhythmic as a function of the CERAD score, while the horizontal axes represent the fGSEA enrichment significance (-Log(BH.q)) of differentially rhythmic genes as a function of Braak score. **(E-G)** Joint pathway analysis using MetaboAnalyst for metabolites and transcripts. **(E)** Pathways cycling in CTL subjects, **(F)** pathways differentially cycling in AD subjects, and **(G)** pathway with differential MESORS comparing AD and control subjects. The vertical axes show pathway significance (-Log(p value)), and the horizontal axes measure pathway impact, which reflects the relative importance of a metabolic pathway based on its degree centrality.

**Figure S7.**
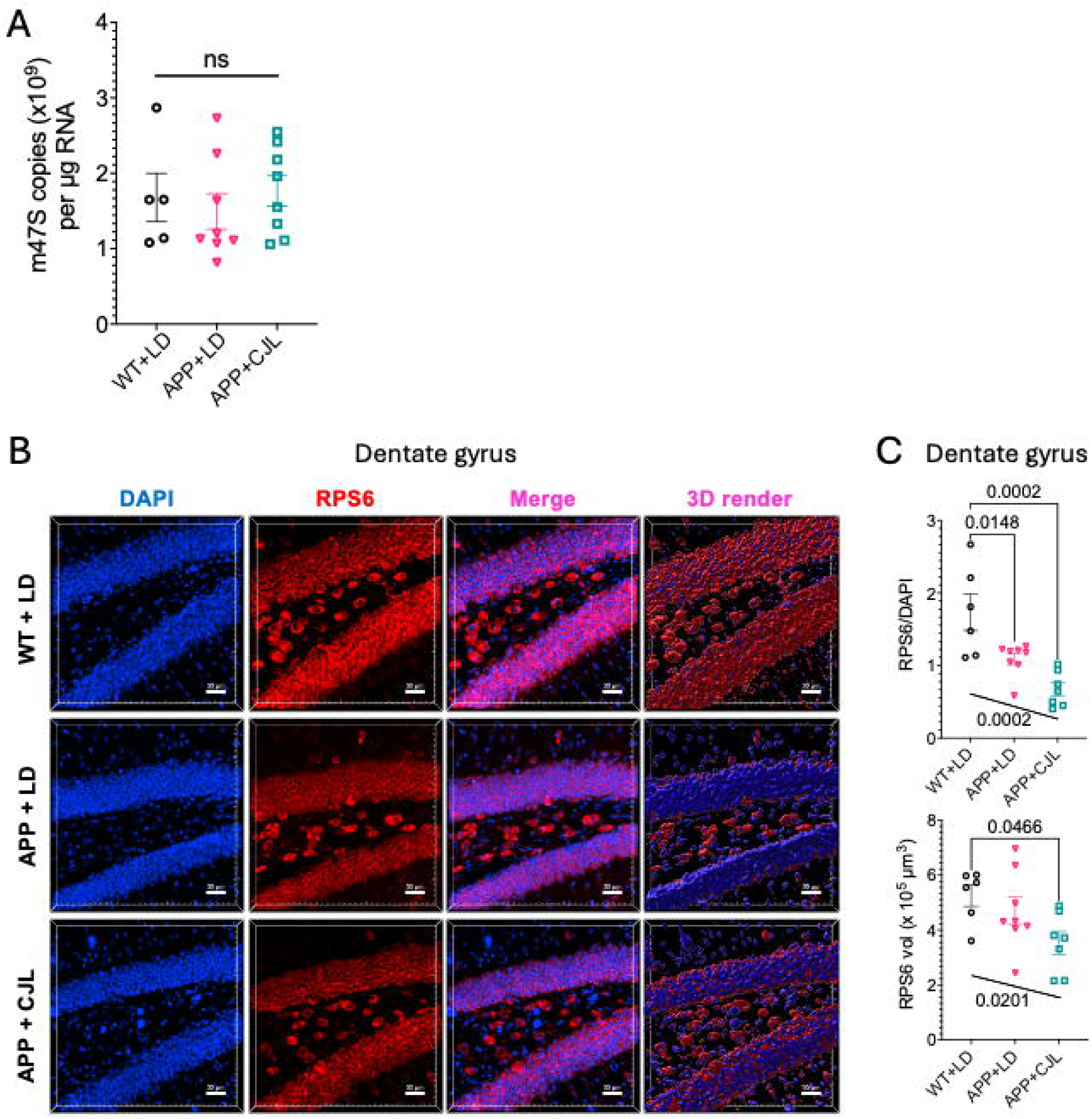
Additional imaging and quantification of ribosomal structural components, related to Figure 6. **(A)** RT-PCR quantification of m47S rRNA in three experimental cohorts. Error bars indicate standard error of the mean (SEM). **(B)** Immunostaining of dentate gyrus slices for RPS6 across experimental groups. **(C)** Quantification of RPS6 immunostaining in dentate gyrus. Error bars indicate standard error of the mean (SEM). Significance values at the top (on horizontal lines) denote Tukey’s multiple comparisons test. Significance values near the bottom, by slanted lines, indicate significance of Kendall’s ranked correlation coefficient.

